# Multiple interactions between Scc1 and Scc2 activate cohesin’s DNA dependent ATPase and replace Pds5 during loading

**DOI:** 10.1101/205914

**Authors:** Naomi J Petela, Thomas G Gligoris, Jean Metson, Byung-Gil Lee, Menelaos Voulgaris, Bin Hu, Sotaro Kikuchi, Christophe Chapard, Wentao Chen, Eeson Rajendra, Madhusudhan Srinivisan, Hongtao Yu, Jan Löwe, Kim A Nasmyth

## Abstract

In addition to sharing with condensin an ability to organize DNA into chromatids, cohesin regulates enhancer-promoter interactions and confers sister chromatid cohesion. Association with chromosomes is regulated by hook-shaped HEAT repeat proteins that Associate With its Kleisin (Scc1) subunit (HAWKs), namely Scc3, Pds5, and Scc2. Unlike Pds5, Scc2 is not a stable cohesin constituent but, as shown here, transiently displaces Pds5 during loading. Scc1 mutations that compromise its interaction with Scc2 adversely affect cohesin’s ATPase activity, loading, and translocation while Scc2 mutations that alter how the ATPase responds to DNA abolish loading despite cohesin’s initial association with loading sites. Lastly, Scc2 mutations that permit loading in the absence of Scc4 increase Scc2’s association with chromosomal cohesin and reduce that of Pds5. We suggest that cohesin switches between two states, one with Pds5 bound to Scc1 that is not able to hydrolyse ATP efficiently but is capable of release from chromosomes and another in which Scc2, transiently replacing Pds5, stimulates the ATP hydrolysis necessary for loading and translocation away from loading sites.

## Introduction

How enhancers activate the correct promoters during development, how chromosomal DNAs are weaved into chromatids, and how sisters are held together during mitosis are all fundamental questions in chromosome biology. These apparently disparate processes are conferred by a pair of related Smc/kleisin complexes called cohesin and condensin. Both contain a pair of rod-shaped Smc proteins that associate to create V-shape heterodimers (Smc1/3 in cohesin) whose ATPases at their apices are bound by the N- and C-terminal domains of a kleisin subunit (Scc1), forming a huge tripartite ring. In addition to conferring sister chromatid cohesion during G2 and M phases (Nasmyth, 2001), cohesin is involved in the process by which the insulator protein CTCF regulates enhancer-promoter interactions (Fudenberg et al., 2016) and is responsible for creating the Topologically Associated Domains (TADs) detected by HiC (Suhas Rao, 2017). Condensin on the other hand is crucial for re-organizing DNA into compact cylindrical chromatids specifically during mitosis (Hirano, 2006).

Two recent findings demonstrate that cohesin and condensin must operate using similar principles. First, cohesin can also organize DNA into chromatids, albeit during interphase (Klein et al., 1999; Tedeschi et al., 2013) and only when its turnover on chromatin is abrogated by inactivation of a regulatory protein called Wapl. Second, cohesin’s association with (Ciosk et al., 2000) and dissociation from (Beckouet et al., 2016) chromosomes are regulated by three related hook-shaped proteins composed of HEAT repeats, namely Pds5, Scc3/SA, and Scc2/Nipbl. All three are monophyletic with equivalent subunits in condensin. This class of regulatory subunit called HAWKs (HEAT repeat proteins Associated With Kleisins) distinguishes cohesin and condensin (Wells et al., 2017) from bacterial Smc/kleisin complexes and the eukaryotic Smc5/6 complex, whose kleisin subunits associate instead with tandem winged helical domain proteins called KITEs (Palecek and Gruber, 2015).

There is substantial evidence that cohesin’s ability to hold sister chromatids together depends on entrapment of sister DNAs inside individual heterotrimeric Smc/kleisin rings (Haering et al., 2008). By analogy, cohesin’s association with chromatin might involve entrapment of individual DNAs (Gligoris et al., 2014). How cohesin and condensin organize DNA into chromatids is not understood, but it has been suggested that they do so by creating small loops of DNA that are then processively extruded from the complex (Nasmyth, 2001), a process now known as loop extrusion (LE). According to a new version of the LE hypothesis, CTCF creates TADs and insulates enhancers from non-cognate promoters by halting the loop extrusion process when cohesin reaches CTCF bound to DNA in a specific orientation (Fudenberg et al., 2016; Sanborn et al., 2015).

The chromodynamics of cohesin are determined by three processes: loading, translocation, and release. Loading is thought to involve entrapment either of individual DNA segments or loops of DNA inside cohesin’s ring. The mechanism of translocation is unclear. It is presumably driven by ATP hydrolysis as recently observed for condensin in vitro (Terakawa et al., 2017). Release involves creation of an exit gate by transient dissociation of the three helical bundle formed between the α ◂ -helix in Scc1’s NTD and the coiled coil emerging from Smc3’s ATPase (Chan et al., 2012; Gligoris et al., 2014). It depends on Pds5 bound to a highly conserved peptide motif (L x L x (D/E) x Ψ x x x (D/E) Φ Φ) situated 10-20 residues C-terminal to α3 (Lee et al., 2016). Release also requires binding of YGR motifs in Wapl’s NTD (Ouyang et al., 2016) to Pds5’s NTD (Chan et al., 2012) as well binding of another part of Wapl to Scc3’s central domain (Beckouet et al., 2016).

If individual DNA segments are entrapped during loading, then the process must involve creation of an entry gate (Gruber et al., 2006). Whether this is the same as the exit gate or is instead located at the Smc1/Smc3 hinge dimerization domain is controversial. Notably, if loading merely involved insertion of DNA loops inside rings, then there would in fact be no need to break any of the ring’s three interfaces. What is clear is that loading depends on Scc2’s hook-shaped C-terminal domain and an unstructured NTD that snakes through a smaller Scc4 subunit composed of a superhelical array of 13 TPR repeats (Ciosk et al., 2000; Hinshaw et al., 2015; Hu et al., 2015). Because Scc3 is required for cohesin’s stable association with chromosomes, it might also be involved in the loading process. In contrast, neither Pds5 (Chan et al., 2013) nor Wapl, which binds to it, are necessary. Crucially, loading requires engagement of Smc1’s and Smc3’s ATPase heads in the presence of ATP as well as the latter’s subsequent hydrolysis (Hu et al., 2011).

Attempts to reproduce loading in vitro have been only partially successful. Stable association with circular DNA of cohesin from *S. pombe* is stimulated by ATP and by its Scc2 ortholog Mis4. Strangely, it is stimulated equally well by Pds5 and Wapl (Murayama and Uhlmann, 2015), neither of which are necessary for loading in vivo. The reaction is also barely affected by mutations such as Smc1E1158Q or Smc3E1155Q that abolish ATP hydrolysis. Imaging studies have documented sliding of cohesin along individual DNAs in vitro but loading in these cases is independent of both ATP and Scc2 (Davidson et al., 2016; Stigler et al., 2016).

Given that the goal of in vitro biochemistry is to reproduce events known to occur in vivo, a key limitation of current in vitro studies is that they have been performed with complexes from organisms in which the in vivo loading process has been documented poorly, if at all. Establishment of bona fide in vitro DNA replication would not have been possible without characterizing events taking place in vivo at known replication origins (Diffley, 2016) and the same applies to cohesin loading. In this regard, the best characterized loading sites are those in the yeast *S. cerevisiae* where there are broadly two populations of chromosomal cohesin complexes: those loaded throughout chromosomes (arm cohesin) and those loaded under the control of their 120 bp point centromeres (*CEN*s), which are responsible for loading the bulk of cohesin that accumulates in peri-centric sequences 30 kb either side of each centromere (Fernius and Marston, 2009; Hu et al., 2011; Weber et al., 2004).

Cohesin appears to translocate into peri-centric sequences soon after loading at *CEN*s and as a consequence, few if any of its subunits accumulate to high levels at *CEN*s themselves. In contrast, Scc2, which is not currently considered a bona fide cohesin subunit but merely a factor required for loading, is concentrated solely at *CEN*s (Hu et al., 2015), presumably because these are sites at which loading takes place at especially high rates. Whether Scc2 accumulates at *CENs* as a component of cohesin complexes undergoing loading or is merely targeted to *CEN*s through association of its Scc4 subunit with inner kinetochore proteins is not known. Remarkably, cohesin complexes containing versions of Smc1 (Smc1E1158Q) or Smc3 (Smc3E1155Q) that can bind but not hydrolyze ATP also associate preferentially at *CEN*s (Hu et al., 2015). Live cell imaging shows that like Scc2, they do so only in a transient manner (Hu et al., 2011). It has therefore been suggested that engagement of cohesin’s ATPase heads in the presence of ATP permits, or indeed actually triggers, cohesin’s association with *CEN* loading sites along with Scc2, but that hydrolysis is required to complete the reaction in a manner that permits translocation into neighbouring chromatin.

Heterozygous null alleles of Scc2’s human ortholog Nipbl are responsible for a large fraction of cases of a multi-organ developmental disorder called Cornelia de Lange syndrome (CdLS) (Krantz et al., 2004), which is thought to be caused by transcriptional defects caused by changes in the dynamics of cohesin’s association with the genome. That Scc2/Nipbl but not Scc1 itself is haplo-insufficient raises the possibility that CdLS might be caused by an altered behaviour of chromosomal cohesin, and not merely by a reduced amount. Nevertheless, explaining this phenomenon clearly requires a better understanding of how Scc2/Nipbl promotes loading and whether it also regulates cohesin’s subsequent translocation.

The experiments described here suggest that cohesin switches between two states: one with Pds5 bound to Scc1 with little or no ATPase activity and a second with greatly elevated ATPase activity due to Pds5’s replacement by Scc2. The importance of this process during loading and translocation is supported by the behaviour of Scc1 and Scc2 mutants that alter the way these two proteins interact. We suggest that Scc2 should no longer be considered merely as a loading factor but as a bona fide cohesin subunit whose replacement of its fellow HAWK Pds5 promotes translocation as well as loading, in both cases through stimulating cohesin’s ATPase activity. Crucially, we demonstrate that among HAWKs, Scc2 alone is both necessary and sufficient for stimulating cohesin’s DNA dependent ATPase activity.

## Results

### Scc2 is necessary and sufficient to stimulate DNA dependent ATPase activity associated with cohesin’s trimeric rings

To address the role of cohesin’s three HAWK subunits in modulating its ATPase, we purified three types of yeast cohesin rings from insect cells: trimers containing Smc1, Smc3, and Scc1; tetramers containing Smc1, Smc3, Scc1, and Scc3; and hexamers containing Smc1, Smc3, Scc1, Scc3, Pds5, and Wapl (Fig. 1A, S1A). Little or no ATPase activity was associated with any of these, even in the presence of DNA (Fig. 1ABCD). However, activity associated with tetramers and hexamers was greatly stimulated by addition of a version of Scc2 whose N-terminal Scc4-binding domain was replaced by GFP (GFP-Scc2) and increased further still by DNA (Fig. 1BC). Importantly, this activity was abolished by Smc1E1155Q Smc3E1158Q double mutations (Smc1/3 EQ; Fig. S1B). In contrast, GFP-Scc2 barely affected activity associated with trimers (in the absence of DNA) but stimulated it upon addition of Scc3 purified from *E. coli* (Fig. 1DE). Thus, at least in the absence of DNA, Scc3 enhances Scc2’s ability to stimulate cohesin’s ATPase. In contrast to Scc2, Pds5 had no effect on the ATPase activity of tetramers, with or without DNA (Fig. S1C). Remarkably, in the presence of DNA, GFP-Scc2 stimulated ATPase activity associated with trimers, to a level comparable to that of tetramers and hexamers treated with GFP-Scc2 (Fig. 1D). These observations imply that among cohesin’s HAWKs, Scc2 is not only necessary for cohesin’s ATPase but also sufficient to confer its responsiveness to DNA. Scc3 clearly enhances ATPase activity, but unlike Scc2 this effect is bypassed by DNA addition. The lack of ATPase activity upon addition of Pds5 and Wapl is striking because it has been suggested that these proteins stimulate loading of *S. pombe* cohesin in vitro in the absence of Scc2 (Murayama and Uhlmann, 2015). Because loading in vivo is independent of Pds5 (see also Fig. 6C) but dependent on Scc2 and abolished by Smc1/3 EQ mutations that abolish ATPase activity, we suggest that Pds5-induced loading may be an in vitro artefact.

**Figure 1.**
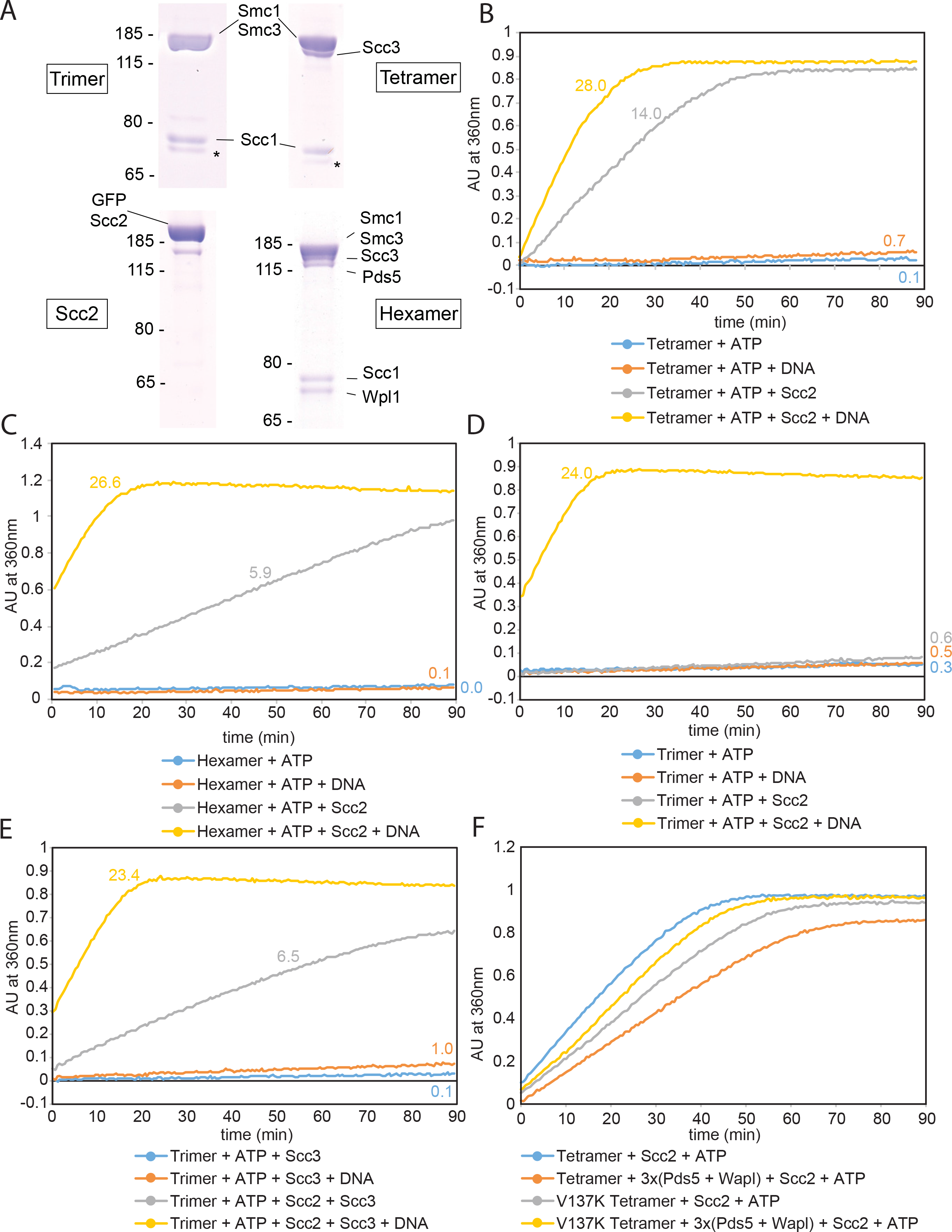
Scc2 drives cohesin’s DNA-dependent ATPase. **(A)** Cohesin trimers (Smc1, Smc3, Scc1), tetramers (Smc1, Smc3, Scc1, Scc3), hexamers (Smc1, Smc3, Scc1, Scc3, Pds5, Wapl) and GFP-Scc2 were affinity purified from Sf9 cell cultures using Strep-trap columns followed by gel filtration. **(B)** Purified hexamers were incubated with DNA, Scc2 or both and the reaction initiated by adding ATP. Rates were calculated by measuring the change in absorption at 360nm over time. **(C)** ATPase activity of tetramers. **(D)** ATPase activity of trimers. **(E)** Effect of Scc3 on ATPase of trimers. **(F)** Effect of three fold excess of Pds5 and Wapl on ATPase of wild type and *scclV137K* tetramers.

Our finding that Scc2 stimulates the ATPase activity of hexamers almost as much as tetramers implies that Scc2 can associate with cohesin and stimulate its ATPase even when the complex was initially occupied by Pds5. Given that Pds5 and Scc2 may compete for occupancy of cohesin, we measured the effect of adding a three-fold molar excess of Pds5 and Wapl to cohesin tetramers. This reduced Scc2-stimulated ATPase activity by 2.5 fold, albeit only in the absence of DNA (Fig. 1F). This inhibition was clearly due to Pds5 binding to cohesin’s kleisin subunit, and not an artefact of merely adding additional protein, because tetramers that cannot bind Pds5 (Scc1V137K) were refractory to inhibition by Pds5/Wapl (Fig. 1F). During the course of these experiments, we noted that *scc1V137K* modestly reduced cohesin’s ATPase, even in the absence of Pds5, implying that the Scc1’s Pds5 binding motif may interact with Scc2 as well as Pds5 (Fig. 1F). Lastly, SDS-PAGE revealed that Pds5 is selectively depleted from cohesin associated with GFP-Scc2 following its addition to wild type or EQ hexamers in the presence of ATP (Fig. S1D).

Our finding that Scc2 has a crucial role in stimulating cohesin’s ATPase activity is consistent with the fact that cohesin’s association with chromosomes is abolished by Smc1/3 EQ mutations and dependent on Scc2. Given that cohesin’s ATPase may also be required for translocation (see below), our findings raise the possibility that Scc2 may regulate the behaviour of cohesin complexes that have already loaded onto chromosomes as well as the loading reaction itself.

### Cohesin associates with Scc2 at *CEN*s and then translocates into peri-centric sequences

The observation that Scc2 interacts with cohesin tetramers or hexamers in vitro raises the question as to whether these interactions occur in vivo, and if so, when. Strangely, ChIP-seq revealed that cohesin tetramers co-localize with Pds5 but not with Scc2 throughout the genome (Hu et al., 2011). Thus, Scc2 does not stably co-localize with cohesin once the latter has loaded onto chromosomes. Crucially, Scc2 can be detected at *CEN*s using calibrated ChIP-seq (Hu et al., 2015) and imaging revealed that it has a 1-2s residence time (Hu et al., 2011). A key question is whether Scc2 associated with *CEN* loading sites is recruited there by the Ctf19 complex (Fernius and Marston, 2009) independently of cohesin or whether Scc2 is instead bound transiently to cohesin rings undergoing loading reactions at *CEN*s.

To address this, we compared the distributions of Scc2 and cohesin’s Scc1 subunit around centromeres using calibrated ChIP-seq in cycling and G1 arrested cells. This revealed that more not less Scc2 accumulates at *CEN*s in α-factor arrested cells than in cycling cells (Fig. 2A). Scc1 is expressed at only low levels in α-factor arrested cells, as indeed are Scc2 and Pds5 (Fig S2B). Despite this, calibrated ChIP-seq reveals that a modest amount of cohesin is associated with peri-centric sequences (Fig. 2C, light blue line), suggesting that loading does in fact occur during this stage of the cell cycle (Hu et al., 2015). Crucially, Scc2’s association with *CEN*s depends on cohesin because it is greatly reduced when cells undergo S phase in the absence of Scc1 (Fig. 2B). The previous conclusion that Scc2 is absent from *CEN*s in pheromone arrested cells as well as Scc1-depleted cells (Fernius et al., 2013) should therefore be revised.

**Figure 2.**
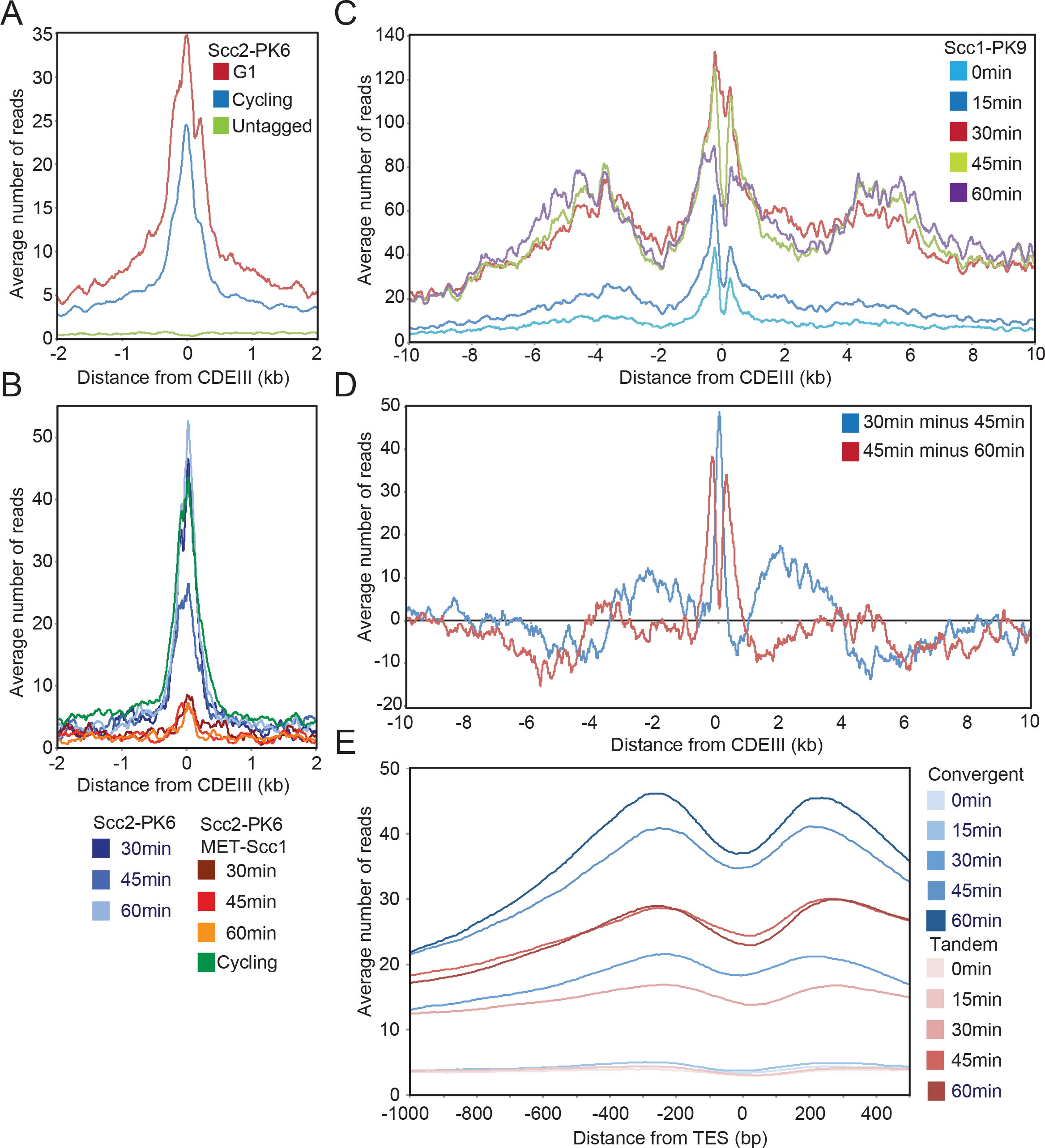
Cohesin associates with Scc2 at *CEN*s and then translocates into pericentric sequences. **(A)** Calibrated ChIP-seq comparing average profile of Scc2-PK6 around *CEN*s in G1 (α factor) and cycling cells. The number of reads at each base pair from *CDEIII* was averaged over all 16 chromosomes. (K21388, K699). **(B)** Average calibrated ChIP-seq profiles of Scc2-PK6 in cells expressing endogenous *SCC1* expressed from the *SCC1* or *MET3* promoter. Cells were arrested in G1 in the presence of methionine prior to release into methionine and nocodazole containing medium. Samples taken at 30, 45 and 60 min after release and from cycling cells grown in the absence of methionine. (K25222, K21388) **(C)** Average calibrated ChIP-seq profiles of Scc1-PK6 every 15 minutes from 0 to 60 min after release from G1. Data reanalyzed from (Hu et al. 2015)(GSE69907). **(D)** Difference plot detailing the difference in average centromere plots between timepoints 30, 45 and 60 min of Fig. 2C. **(E)** Average calibrated ChIP-seq profiles of Scc1-PK6 around TES of genes longer than 2kb. Profiles of convergent and tandem genes are compared every 15 minutes from 0 to 60 min after release from G1. Data reanalyzed from (Hu et al. 2015)(GSE69907). See also Fig. S2.

The dependence of most peri-centric cohesin on *CEN*s and the inter-dependence of Scc1 and Scc2’s association with these sites suggests that most peri-centric cohesin complexes are derived from those loaded at *CEN*s in a reaction involving Scc2’s transient association with cohesin. Consistent with this notion, re-plotting previously published data (Hu et al., 2015) reveals that in late G1 newly synthesized Scc1 associates initially in a peak centered on *CEN*s and subsequently translocates to neighboring peri-centric regions (Fig. 2C). Thus, a plot of the difference between cohesin’s calibrated ChIP-profile at 30 and 45 min, as well as 45 and 60 min, following release from pheromone reveals that a net movement of cohesin away from *CEN*s themselves and from broad peaks about 2 kb either side of them accompanies its accumulation further away in broad peaks 5 kb either side of *CEN*s (Fig. 2D).

### Cohesin loaded on chromosome arms translocates from gene bodies to 3’ ends

If Scc2’s preferential association with cohesin engaged in loading were a general phenomenon then its ChIP-seq profile might also reveal where cohesin loads along chromosome arms. To address this, we plotted average values of Scc2 after aligning all genes around their transcription start or termination sites (TSS or TES, respectively). The average number of reads for PolII genes, whether from cycling or pheromone arrested cells, were much lower than cohesin and merely 3 fold above the untagged control. Chromosomal Scc2, as measured in this manner, was preferentially excluded from both TSSs and TESs but otherwise did not vary greatly throughout genes (Fig. S2EF). Though it was not greatly enriched on ribosomal protein genes or indeed on their promoters (Fig. S2G), higher levels were detected close to the start sites of tRNA genes (Fig. S2H), co-localizing with Smc3E1155Q containing cohesin complexes and adjacent to a peak of wild type cohesin (Fig. S2I). If these profiles reflect cohesin in the process of loading, which is uncertain, then loading must occur fairly uniformly throughout transcription units. The suggestion that cohesin might load preferentially at tRNA genes is more interesting as a tRNA gene has been implicated in generating cohesion in the vicinity of the silent mating type locus *HMRa* (Dubey and Gartenberg, 2007).

Given the inconclusive nature of the above experiments, we re-analysed calibrated Scc1 ChIP-seq profiles from cells released from a G1 arrest induced by α-factor (Fig. 2E). This revealed that soon after loading in late G1, cohesin is distributed uniformly across transcription units and only accumulates at the 3’ end of genes, especially convergent ones, as cells undergo S phase, an event accompanied by Smc3 acetylation and reduced turnover (Fig 2E). Consistent with the notion that cohesin accumulates at the 3’ end of convergent genes only after longer periods of association, inactivation of Wapl accentuated accumulation at the 3’ end of genes in cells blocked in late G1 by the Cdk1 inhibitor Sic1 (Fig. S2J). The simplest explanation for our data is that loading occurs throughout transcription units and not specifically at TSSs. Contrary to previous claims, cohesin does not strictly speaking accumulate between convergent transcription units (Filipski and Mucha, 2002; Lengronne et al., 2004) but rather as a bimodal peak on either side of the TES.

### Scc2 replaces Pds5 during loading at *CEN*s

Accumulation at *CEN*s of cohesin complexes that bind but do not hydrolyze ATP, namely those containing Smc3E1155Q or Smc1E1158Q (Hu et al., 2011; Hu et al., 2015), implies that ATP hydrolysis is not required for cohesin’s association with *CEN* loading sites but instead for a manner of association that permits translocation. Because accumulation at *CEN*s is abolished by *smc1S1130R* or *smc3S1128R* mutations, which prevent ATPase head engagement, it is thought that loading can be broken down into two steps. First, ATPase head engagement promotes co-localization of Scc2 and cohesin at *CEN*s while, second, ATP hydrolysis triggers stable association with and translocation along chromatin.

If Scc2 actually becomes part of the cohesin complex during the first step as opposed to merely co-localizing on the chromosome, then expression of *smc3E1155Q* or *smc1E1158Q* from ectopic genes (endogenous ones are kept intact) should increase Scc2 associated with *CEN*s. Note that in these experiments the calibrated ChIP-seq profiles are therefore a composite of wild type and EQ mutant cohesin. Fig. 3A shows that *smc3E1155Q* expression increases Scc2’s association with *CEN*s at least tenfold but had little effect elsewhere in the genome. It also greatly increased Scc1 at *CEN*s but had little effect elsewhere (Fig. 3B). Scc3’s *CEN* recruitment was also elevated by *smc3E1155Q* (Fig. 3C), albeit more modestly. In contrast, *smc3E1155Q* had little or no effect on the distribution of Pds5 (Fig. 3D), implying that this particular regulatory subunit is absent from *CEN*-associated Scc1/Smc1/Smc3E1155Q/Scc2/Scc3 complexes. Similar results were obtained in cells expressing *smc1E1158Q* (Fig. S3ABCD). Scc3’s presence and Pds5’s absence from such complexes is consistent with the finding that accumulation of GFP-tagged Smc3E1155Q at centromeres depends on Scc3 but not on Pds5 (Hu et al., 2011).

**Figure 3.**
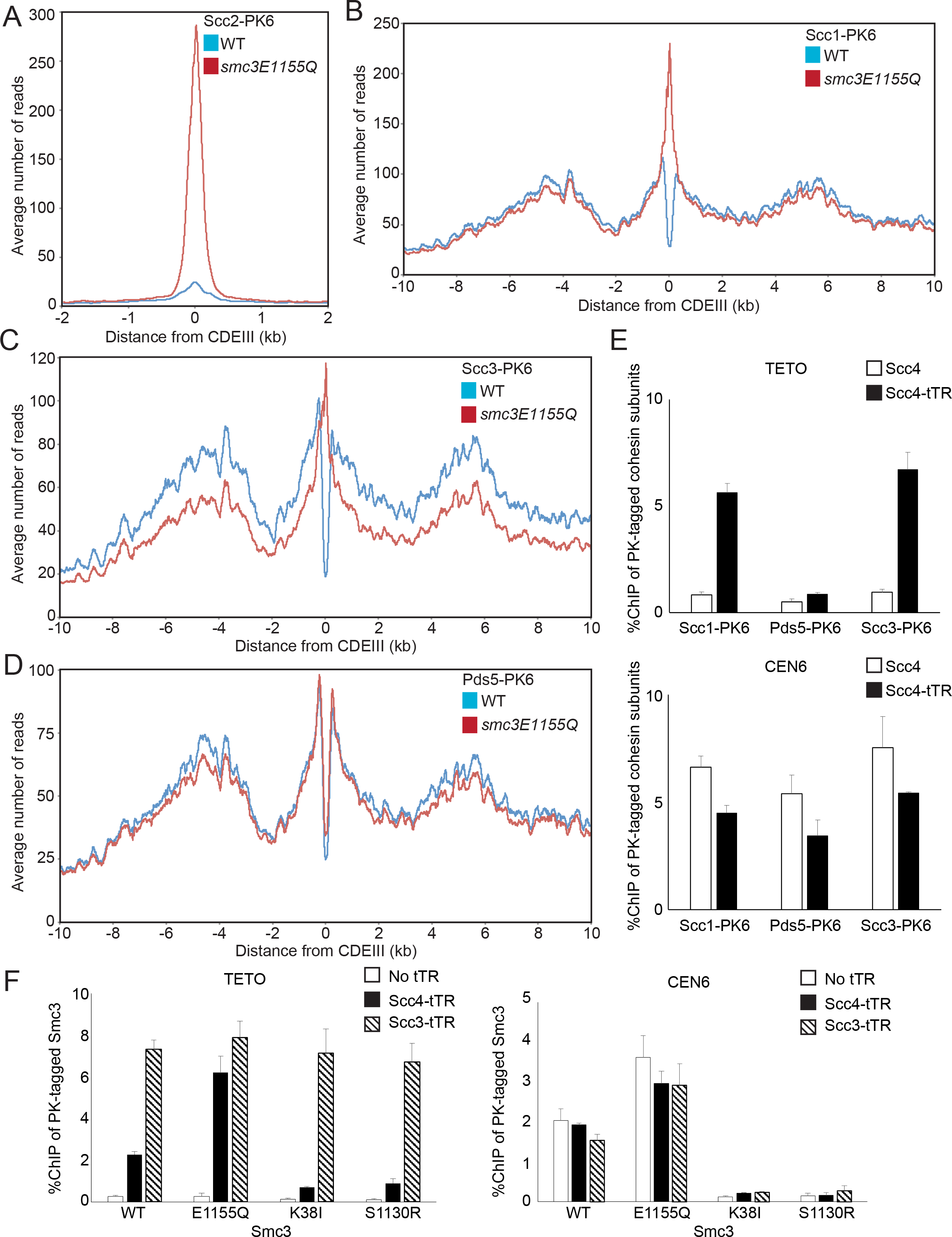
Scc2 replaces Pds5 during loading at *CEN*s. Average calibrated ChIP-seq profiles around *CEN*s comparing localization of **(A)** Scc2, **(B)** Scc1, **(C)**, Scc3 and **(D)** Pds5 in the presence or absence of *smc3(E1155Q)* with WT in cycling cells (K25467, K21388, K25370, K22005, K25373, K17438, K25376, K19012). **(E)** Calibrated qPCR ChIP was used to measure association of PK-tagged Scc1, Pds5, or Scc3 with TETO on chromosome X and a sequence 400 bp from *CEN6* in Scc4-tTR/TETO cells expressing PK-tagged Scc1 (B1674), Pds5 (B1665), or Scc3 (B1625) grown to log phase. Cells with untagged Scc4 (B1627, B1635, and B1667) were used as controls. Each data represents the average of three replicates and S.D. is indicated. **(F)** The tTR-tagged Scc4 or Scc3/TETO diploid cells in which one of two *SMC3* alleles is fused with PK6 tag and mutated (B1612/B1795, B1684/B1796, B1749/B1797, and B1751/B1798) were grown at 25°C. Association of PK-tagged Smc3 mutants with TETO and centromere loci was measured by calibrated ChIP-qPCR. Cells without tTR tag (B1664, B1685, B1748, and B1750) were used as controls.

To address whether Pds5 is excluded from wild type cohesin engaged in loading, we compared the chromosomal profiles of Scc1 and Pds5 in cells arrested in late G1 when Scc1 is expressed at high levels but cohesin associated with peri-centric sequences is known to be turning over rapidly (Chan et al., 2012) (Fig. S3E). A scatter plot shows that Scc1 and Pds5 levels correlate highly throughout the genome. However, there is a set of sequences whose slope is half the average, namely sequences selectively depleted of Pds5, which correspond to *CEN* sequences (Fig. S3F). Depletion of Pds5 from *CEN*s was less pronounced in G2/M phase where loading is less active (Fig. S3GH). Pds5 is therefore displaced by Scc2 at *CEN*s even in wild type. The profiles suggest that Pds5 re-associates with cohesin and replaces Scc2 by the time the complex has translocated approximately 300bp from the loading site.

Further evidence that Scc2’s association with cohesin in vivo is associated with displacement of Pds5 is our finding that tethering Scc4 to Tet operators on chromosome X recruits Scc1 and Scc3 but very little Pds5 to this locus (Fig. 3E). Interestingly, recruitment of Smc3 to Tet operators bound by Scc4 was increased by Smc3E1155Q but greatly reduced by Smc3K38I or Smc3S1130R, implying that Scc2 interacts preferentially with cohesin whose ATPase heads are engaged in the presence of ATP (Fig. 3F). In contrast, these Smc3 mutations had little or no effect on Smc3’s recruitment to Tet operators by Scc3-TetR, suggesting that binding of Scc2 may be uniquely sensitive to the state of ATPase head engagement.

Our finding that Scc2’s association with cohesin is accompanied by loss of Pds5 and that Scc2 but not Pds5 stimulates cohesin’s ATPase suggests that it is the Scc2-bound version that is capable of loading and translocation. If so, Pds5 should be unnecessary for these processes. As predicted, Pds5 depletion had no adverse effect on either loading or translocation of cohesin, at least in late G1 cells (Fig. 6C), where Pds5’s role in promoting Smc3 acetylation (Chan et al., 2013) would be immaterial. Because both Smc3E1155Q and Smc1E1158Q greatly increase the amount of Scc1 and Scc2 associated with cohesin at *CENs,* we suggest that ATP hydrolysis is normally necessary for Scc2’s subsequent replacement by Pds5, an event that occurs to most chromosomal cohesin complexes in yeast.

In summary, the loading process at *CEN*s can be divided into two major steps. During the first, engagement of ATPase heads in the presence of ATP is accompanied by replacement of Pds5 by Scc2. During the second, ATP hydrolysis completes the reaction, leads to cohesin’s stable association with chromatin and Scc2’s replacement by Pds5. Our experiments do not address whether further rounds of ATP hydrolysis mediated by the Scc2-bound form of cohesin promote translocation into peri-centric sequences.

### Scc2 residues involved in DNA-dependent ATPase activity are required for the late loading step

The finding that association of Smc3E1155Q with centromeres depends on Scc2 suggests that Scc2 is required for the first step (Hu et al., 2011). Is it also required for the second? With the aim of identifying mutations that might be preferentially defective in the second step, we created a series of mutations in highly conserved Scc2 surface residues (Fig. S4A) as well as some residues mutated in CdLS patients (Table S1). Scc2 exhibits a ribbon of conservation that twists in a complex manner around the hook-shaped protein (Kikuchi et al., 2016) (Fig S4A). Remarkably, no single surface amino acid change in the untagged endogenous locus was found to be lethal (Table S1) but two double mutants were, namely *S717L K721E* and *K788E H789E* (Fig. 4AB and Fig. S4C). Both affect the part of Scc2 that is most conserved among HAWK subunits, the region composed of canonical HEAT repeats that unlike the rest of the protein are not twisted. The residues equivalent to K788 and H789 are invariably basic in a very wide variety of eukaryotes and might therefore have a role in binding DNA.

**Figure 4.**
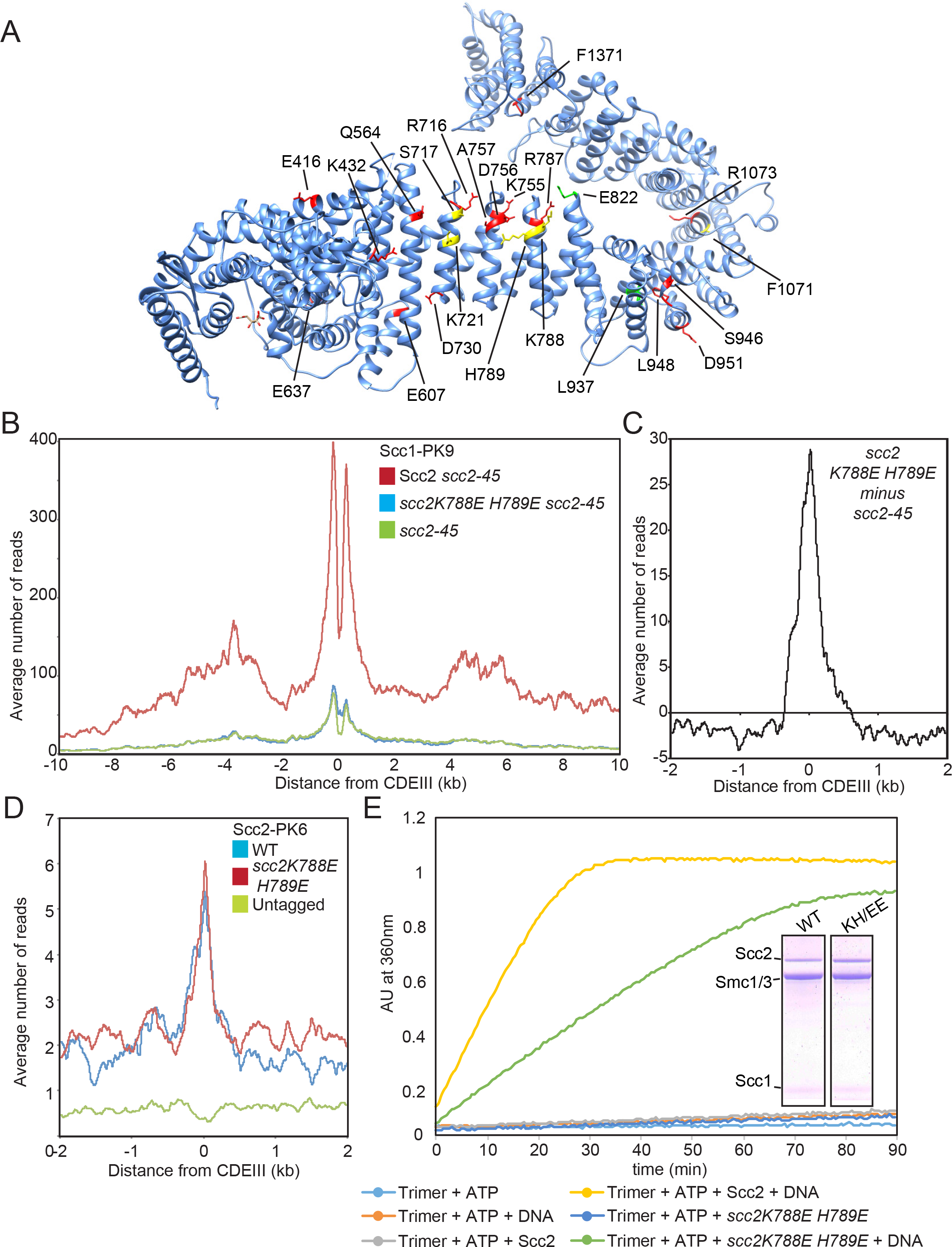
Scc2 is required for both early and late loading steps. **(A)** Residues mutated in *S. cerevisiae* Scc2 mapped onto the *C. thermophilum* structure (PDB 5T8V). Lethal single or double mutations in yellow, viable mutations in red, and gain of function mutations in green. **(B)** Average calibrated ChIP-seq profiles of Scc1-PK9. Cells expressing WT, *K788E H789E* double mutant or no ectopic copy of *SCC2* over endogenous *scc2-45* were arrested in G1 at 25°C before release at 37°C into medium containing nocodazole. Samples were taken 75 min after release. (K24188, K24185, K22390) **(C)** The average profile of *scc2-45* was subtracted from that of *scc2E822K L937Fscc2-45,* producing a difference plot revealing loading due to *scc2E822KL937F.* **(D)** Average calibrated ChIP-seq profiles of ectopic WT and *scc2K788E H789E* in cycling cells (K25185, K25186, K699) **(E)** ATPase of WT trimers in the presence or absence of WT or mutant Scc2, and DNA. A fraction of the mix was analysed by Coomassie staining following SDS-PAGE to confirm protein levels. See also Fig. S4.

To evaluate the effect on loading, we expressed either wild type or *scc2K788E H789E* from an ectopic locus in cells harbouring the thermosensitive (ts) *scc2-45* (*L545P D575G*) allele. Unlike wild type, *scc2K788EH789E* failed to suppress the genome-wide loading defect of *scc2-45* cells (Fig. 4B). Remarkably, it did support Scc1’s association with *CEN* loading sites (Fig. 4C). Indeed, calibrated ChIP-seq revealed that the mutant protein associates with *CEN*s as efficiently as wild type (Fig. 4D), as did live imaging of GFP-tagged proteins (Fig. S4D). These data suggest that Scc2K788E H789E can support the first step in the loading reaction, namely association with cohesin at *CEN*s, but cannot support stable association (loading itself) or translocation into neighbouring sequences. Because wild type is capable of both steps, it is difficult to evaluate whether the mutant is as efficient as wild type in completing the first step.

We conclude that Scc2 is required not only for association of cohesin with engaged ATPase heads at *CEN* loading sites but also for converting these into complexes that stably associate with chromatin and are capable of translocating along it. *scc2K788E H789E* reduced by about two fold Scc2’s ability to stimulate the ATPase activity of cohesin tetramers in vitro, even in the presence of DNA (Fig. S4F). The defective response to DNA caused by *scc2K788E H789E* is especially apparent with trimers, which fail to respond to Scc2 unless DNA is added. Stimulation of trimer ATPase by DNA was halved by *scc2K788E H789E* (Fig. 4E). Thus, K788 and H789 may indeed have an important role in the mechanism by which DNA stimulates ATPase activity associated with Scc2-bound cohesin. A crystal structure of condensin’s Ycg1 HAWK subunit associated with both DNA and the γ-kleisin Brn1 reveals that Ycg1 adopts a structure more similar to that of Scc2 than Scc3 (Kschonsak et al., 2017), and that a highly conserved R253 residue contacting phosphates on the DNA backbone corresponds to Scc2’s K788 (Fig. S4B). A role in contacting DNA might therefore be a feature conserved between Scc2 and Ycg1.

### A gain-of-function *SCC2* allele (*scc2E822KL937F*) that bypasses Scc4 for loading on arms but not at *CEN*s

Loading of cohesin at *CEN*s as well as along chromosome arms depends on Scc4 bound to Scc2’s NTD (Fig. 5A). By selecting revertants of the ts allele *scc4-4* (Y40N) capable of proliferation at 37°C, we identified two *SCC2* alleles that permitted proliferation without *SCC4.* Tetrad analysis revealed that *scc2E822K* was a better suppressor than *scc2L937F* and the *scc2E822KL937F* double mutant better still (Fig. S4G). E822 is a highly conserved surface residue situated on the spine of the Scc2’s most conserved HEAT repeats, very close to K788 H789 (Fig. 4A, S4A, S5A). L937 is invariably a hydrophobic residue and is buried within a HEAT repeat α-helix close to the point where the protein starts to bend back on itself. Substitution by phenylalanine presumably alters the way the helix interacts with its neighbour and bulky aromatic residues are rarely if ever found at this position. *scc2E822K L937F* enhances cohesin’s association with chromosome arms (Fig. S5D) and causes it to accumulate to higher than normal levels in two peaks 500 bp on either side of *CENs,* suggesting that it may retard translocation (Fig. 5B, S5D). The double mutation elevates loading along chromosome arms in ts *scc4-4* mutants from ~25% to ~80% of wild type when cells undergo S phase at 37°C in the presence of nocodazole, but barely suppresses the loading defect at *CEN*s (Fig. 5AB). *scc2E822K L937F* increased modestly Scc2’s ability to stimulate ATPase activity associated with cohesin tetramers (Fig. 5C).

**Figure 5.**
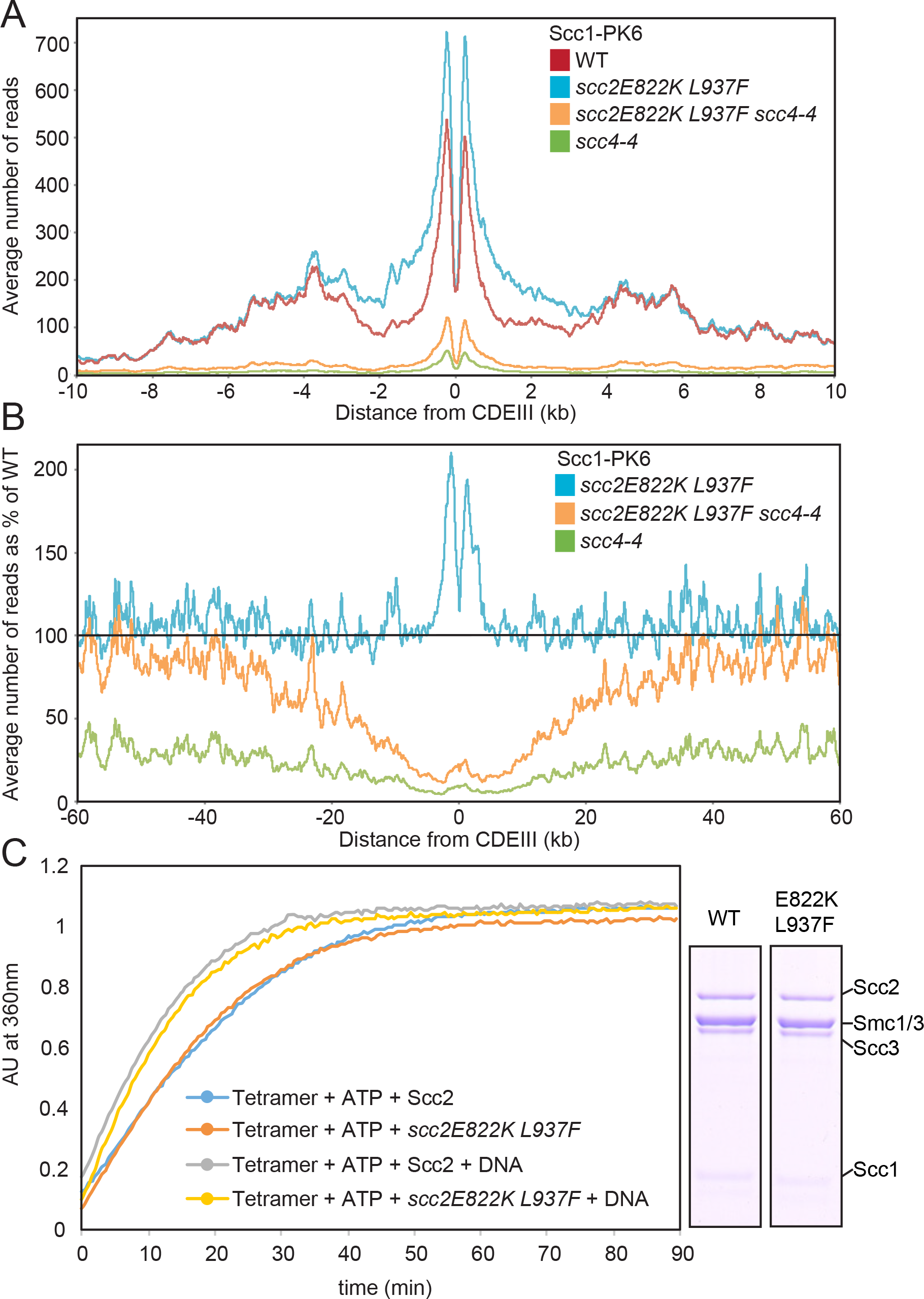
*scc2E822K L937F* bypasses Scc4 on arms but not at *CEN*s. **(A)** Average calibrated ChIP-seq profiles of Scc1-PK6 in *SCC2 SCC4, scc2E822K L937F SCC4, SCC2 scc4-4,* and *scc2E822KL937F scc4-4* cells arrested in G1 at 25°C before release at 37°C into medium containing nocodazole. Samples were taken 75 min after release (K22005, K24687, K24744, K22001). **(B)** Average calibrated ChIP profiles 60 kb either side of *CDEIII* plotted as a percentage of the average number of reads obtained for WT *SCC2 SCC4* cells in A. **(C)** Effect of *scc2E822K L937F* on tetramer ATPase. A fraction of the reaction analysed by Coomassie staining following SDS-PAGE. Fig. S5 shows the effect of *scc2E822K L937F* at 25°C, where its enhancement of loading is greater than at 37°C.

**Figure 6.**
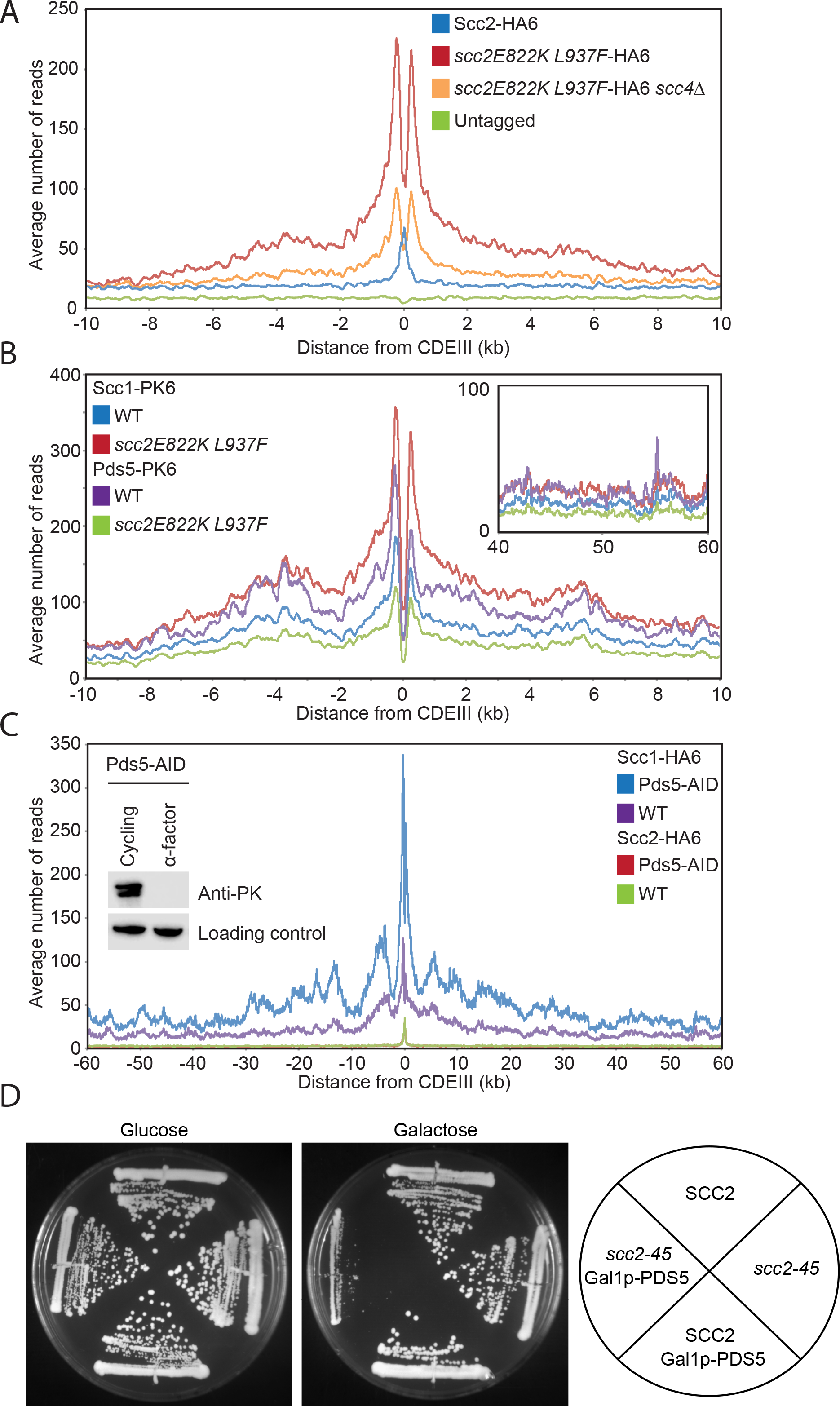
competition between Scc2 and Pds5 regulates cohesin loading genome wide. **(A)** Average calibrated ChIP-seq profiles of WT Scc2-HA6 and *scc2E822K L937F-* HA6 in presence or absence of *SCC4* in cycling cells (K25054, K25053, K25418, K699). **(B)** Average calibrated ChIP-seq profiles of Scc1-PK6 or Pds5-PK6 in the presence of WT or *scc2E822K L937F* in *GAL-SIC1* arrested cells (K26292, K26244, K25999, K25988). **(C)** Average calibrated ChIP-seq profiles of Scc1-HA6 or Scc2-HA6 in the presence or absence of Pds5. Cells were arrested in late G1 using *GAL-SIC1* after addition of 5mM auxin 30 min prior to release from pheromone (K26270, K26277, K26273, K26274). **(D)** Pds5 overexpression hampers growth of *scc2-45* cells. WT (K699), *scc2-45* (B443), *GAL-PDS5* (B1289), and *scc2-45 GAL-PDS5* (B1282) were grown on YEP plates containing glucose or galactose at 30°C for 2 days. See Fig. S5 for FACs profiles.

### *SCC4* is essential for loading cohesin at *CEN*s in the absence of spindle poisons but not in their presence

We subsequently established that *scc4Δ* abolishes loading dependent on the Ctf19 complex subunit Chl4 in cycling *scc2E822KL937F* cells (Fig. S5E), suggesting that Scc4 has a specific role in loading at *CEN*s that cannot be bypassed by *scc2E822KL937*. This is consistent with the recent finding that phosphorylation of Ctf19 by DDK creates a binding site for Scc4, which presumably directs Scc2/4 to *CEN*s and promotes *CEN*-specific loading (Hinshaw et al., 2017). Surprisingly, *scc2E822KL937F* permitted substantial loading around *CEN*s in *scc4Δ* mutant cells arrested in G2/M by nocodazole (Fig. S5E). Crucially this loading also depended on *CHL4,* confirming that it is driven by *CEN* activity. Thus, in addition to Scc4-dependent loading at *CEN*s in cycling cells, an Scc4-independent mechanism promotes loading in the presence of nocodazole.

Because of these findings, we re-investigated the behaviour of *scc4^*m35*^* (F324A K327A K331A K541A K542A), which has been claimed to abolish *CEN*-specific loading in nocodazole arrested cells (Hinshaw et al., 2015). We discovered that the residues mutated in *scc4^*m35*^* are in fact only conserved in organisms closely related to *S. cerevisiae* and *K. lactis*, in other words in yeasts that possess point centromeres (Fig. S5F). Contrary to a previous report, *scc4^*m35*^* did not abolish *CEN*-specific loading when cells are treated with nocodazole, but did so in cycling cells (Fig. S5GH). Our data suggest that the patch altered in *scc4^*m35*^* may be essential for recruitment of Scc4 to *CEN*s and that this process is crucial for *CEN*-specific loading in cycling cells. Our findings suggest that recruitment of Scc4, and thereby Scc2, to *CEN*s by the Ctf19 complex co-evolved with point centromeres.

### Scc2E822K L937F persists on cohesin and displaces Pds5 after loading and translocation

*scc2E822KL937F*has a striking effect on Scc2’s distribution around *CEN*s. Instead of a narrow peak centred on *CDEIII*, the mutant protein accumulated throughout a broad peri-centric interval and especially so with a pair of symmetrical peaks on either side of *CEN*s (Fig. 6A), in a manner reminiscent of cohesin itself in such cells (Fig. 5A). Importantly, *scc4Δ* greatly reduced association of Scc2E822K L937F with peri-centric sequences implying that this association is *CEN*-specific (Fig. 6A). This is accompanied by a dramatic re-distribution of Pds5. Whereas *scc2E822K L937F* causes a modest increase in the amount of Scc1 associated with peri-centric sequences, it causes a 2.5 fold decrease in the amount of Pds5 (Fig. 6B). The net effect is that occupancy of chromosomal cohesin by Pds5 is reduced about three fold around *CEN*s. *scc2E822KL937F* also reduced Pds5’s occupancy of cohesin along chromosome arms by about two fold (*Fig. 6B*). These observations demonstrate that Scc2 displaces Pds5 not only during the process of loading at *CEN*s but also during or after translocation into peri-centric sequences. Our observations suggest that Scc2E822K L937F competes with Pds5 on chromosomal cohesin more effectively than the wild type protein.

### Pds5 inhibits cohesin loading genome-wide

Our finding that Scc2’s occupancy of chromosomal cohesin is accompanied by displacement of Pds5 suggests that the latter might act as a negative regulator of cohesin activities mediated by Scc2, namely loading and possibly also translocation. To address this, we investigated the effect of depleting Pds5 on the distributions of Scc1 and Scc2. To avoid complications associated with the fact that Pds5 is necessary for Smc3 acetylation during S phase, we analysed the effect in cells blocked in late G1 by the Cdk1 inhibitor Sic1. Though Pds5 depletion using an AID degron had little or no effect on Scc2’s distribution, it had a major effect on Scc1, increasing loading throughout the genome twofold (Fig. 6C).

To address whether this effect is an indirect consequence of compromising recruitment of Wapl, whose association with Pds5 is required for cohesin turnover, we also analysed the effect of Wapl deletion (*wpl1Δ*) at this stage of the cell cycle. Interestingly, *wpl1Δ* caused a major increase in peri-centric cohesin, an effect that is probably not due to decreased turnover, because *scc3K404E* (Beckouet et al., 2016) had little effect (Fig. S6A). Importantly, *wpl1Δ* had little or no effect on the extent of Scc1’s association with chromosome arms, implying that the effect of Pds5 depletion is due to increased loading not reduced turnover. We suggest therefore that Pds5 negatively regulates cohesin loading mediated by Scc2 throughout the genome. Consistent with this notion, over-expression of Pds5 from the *GAL* promoter causes lethality in *scc2-45* cells growing at the permissive temperature (Fig. 6D). We do not understand why Pds5’s depletion does not increase Scc2’s association with the genome but suspect that, even in the absence of Pds5, Scc2’s turnover on chromosomal cohesin complexes remains too rapid for efficient formaldehyde fixation. The negative effect of Pds5 on cohesin loading genome-wide in G1 cells is consistent with our finding that Pds5 and Wapl reduce Scc2’s ability to stimulate ATPase activity associated with cohesin tetramers (Fig. 1E).

### Scc2 promotes loading, translocation, and ATPase activity by interacting with multiple Scc1 motifs

To evaluate the importance of Scc2’s association with cohesin, we investigated its mechanism. It has been suggested on the basis of peptide arrays that Mis4 (Scc2) from *S. pombe* functions by binding to Scc3 and to the coiled coils of Smc1 and Smc3 (Murayama and Uhlmann, 2014). In contrast, Scc2 from *C. thermotolerans* binds exclusively to Scc1, to sequences between 126 and 230 (Kikuchi et al., 2016). The mode of binding in *S. cerevisiae* may be similar as Scc2 co-precipitated with a fragment of an N-terminal fragment yeast Scc1 containing residues 1-566 (Fig. 7A). Nevertheless, there is hitherto no evidence that the Scc1:Scc2 interaction is of physiological importance, especially as many of the mutations that supposedly compromise association of *Chaetomium* Scc2 with Scc1 have no phenotype in yeast (e.g. *C. thermotolerans* Scc2 K1018E, R1053Q, R1090T; Table S1) or, more worrying, involve substitutions to residues that are frequently found in other fungi (e.g. *C. thermotolerans* Scc2 L1373P).

**Figure 7.**
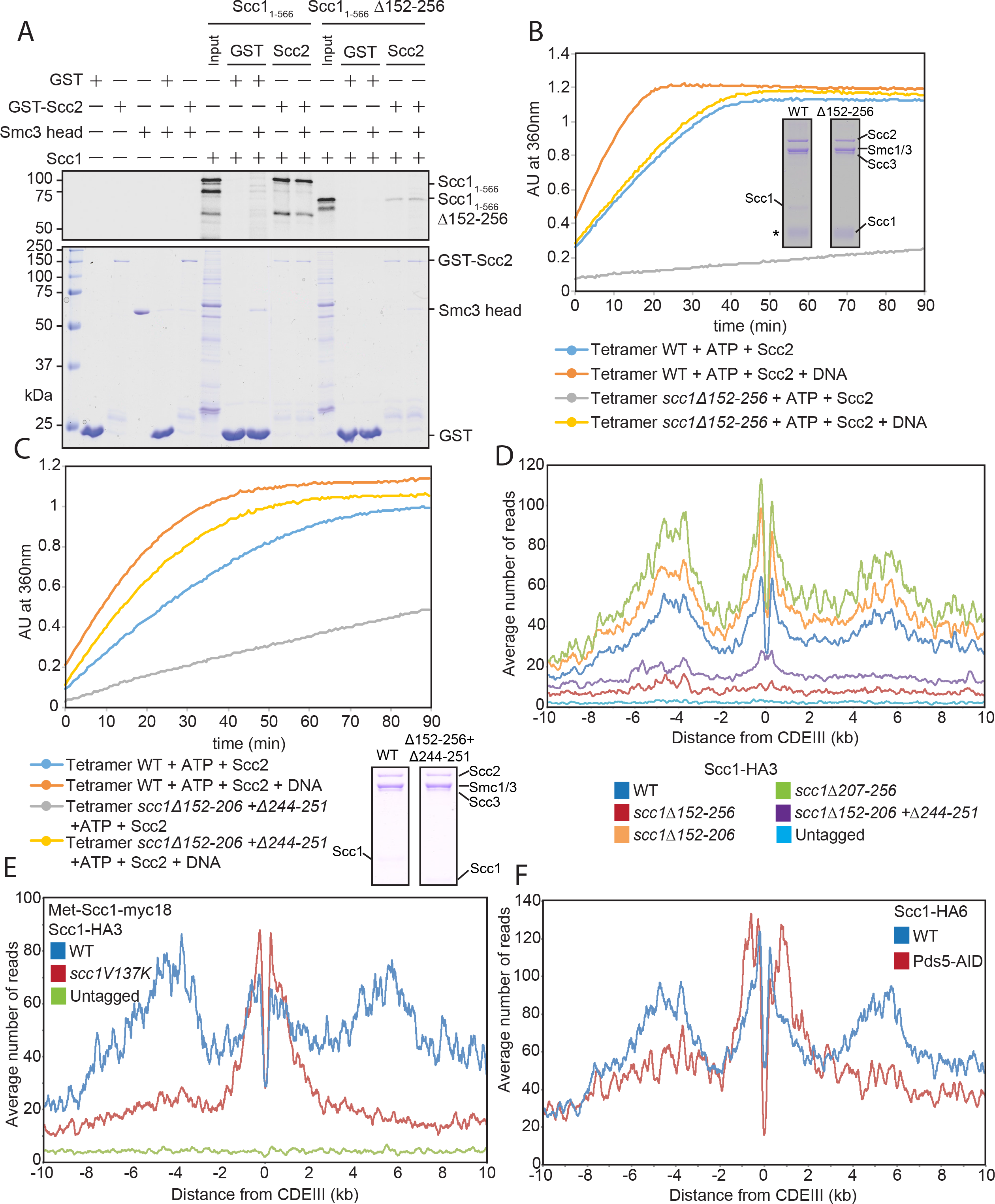
Scc2 promotes loading, translocation, and ATPase activity by interacting with multiple Sccl motifs. **(A)** GST or GST-Scc_2171-1504_ proteins were immobilized on Glutathione Sepharose beads. Beads were incubated with ^35^S-labeled WT or Δ152-256 Scc1_1-566_ in the presence or absence of Smc3head. Input and bound proteins were separated by SDS-PAGE and stained with Coomassie (Bottom) and analysed with a phosphorimager (Top). **(B)** Effect of *scc1Δ152-256* on tetramer ATPase activity. A fraction of the mix was then analysed by Coomassie staining following SDS-PAGE. *The Scc1Δ152-256 protein is masked by purine nucleoside phosphorylase (PNP). **(C)** Effect of *scc1Δ152-206 Δ244-251* on tetramer ATPase. **(D)** Average calibrated ChIP-seq profiles of HA-tagged WT and mutated Scci proteins in cycling cells at 25°C in the presence of untagged *SCC1* (K17184, K25896, K26660, K25995, K25997, K699). **(E)** Average calibrated ChIP-seq profiles of Scc1-HA3 in cells expressing endogenous *SCC1* under either the *MET3* promoter or the endogenous *SCC1* promoter. Cells were arrested in G1 (α factor) in the presence of methionine prior to release into methionine containing medium. Samples were taken at 60 min after release (K25222, K26248, K26251). **(F)** Average calibrated ChIP-seq profiles of Scc1-HA6 in the presence or absence of Pds5. Cells were arrested in G1 by pheromone prior to release for 90 min into auxin and nocodazole containing medium (K26270, K26277). See also Figs. S6 and S7.

Reasoning that Scc1 sequences bound by Scc2 have a role in cohesin loading, we measured the effect on yeast cell proliferation and cohesin loading of deleting Scc1 sequences other than those already known to bind Pds5 (*S.c.* Scc1 131-138), Scc3 (319-393), Smc3 (1-104), and Smc1 (483-566) (Fig. S6C). Tetrad analysis showed that when expressed from an ectopic locus, all such deletions removing no more than 55 amino acids complemented *scc1Δ* (Fig. S6B). However, larger deletions were not able to do so. Thus, while *scc1Δ152-206* and *Δ207-256* conferred viability, *Δ152-256* could not, despite binding Scc3 and forming trimeric rings with Smc1/3 (Fig. S6D, S7B). Crucially, *Δ152-256* caused a severe loading defect throughout the genome (when measured in cycling cells expressing untagged wild type protein (Fig. 7D)), reduced Scc2-dependent ATP hydrolysis (Fig. 7B), especially in the absence of DNA, and reduced binding of Scc1 1-566 to GST-Scc2 in vitro (Fig. 7A).

The loading defect associated with *Δ152-256* could be due either to a length requirement or redundancy of sequence motifs. In the case of the latter, it should be possible to identify within the *207-256* interval a motif whose deletion is lethal when combined with *Δ152-206*. Despite poor conservation among ascomycetes, we noticed that sequences within the interval 244-251 are conserved among yeasts with point centromeres, with a consensus DWDLGITE (Fig. S6F). Indeed, the double mutant *Δ152-206 Δ244-251* was lethal, reduced ATPase activity in vitro (Fig. 7C), as well as loading in vivo (Fig. 7D). Because *Δ180-256* is also lethal, the 180-206 interval must likewise contain an element that is essential when sequences between 207 and 256 are deleted. Interestingly, this interval contains a motif with the consensus LDLDFD, which resembles the 244-251 sequence. We note that a sequence with similar properties (consensus LDLELDFGEDID) is conserved among ascomycetes related to *C. thermotolerans*, within the 126-200 interval known to bind Scc2. *Δ196-203* is viable but its combination with *Δ244-251* is lethal (Fig. S6BC). We therefore suggest that Scc2 promotes cohesin loading in yeast via these two DL motifs.

These motifs cannot be the sole means by which Scc2 interacts with cohesin because in the presence of DNA, Scc2 stimulates the ATPase activity of tetramers containing *scc1Δ152-256* (Fig. 7B). Indeed, GFP-Scc2 binds to mutant tetramers in vitro, albeit slightly less efficiently than wild type (Fig. S7B). Given that Pds5 and Scc2 appear to compete for binding to cohesin in vivo, we considered that Scc2 might also interact with the Pds5-binding motif approximately 20 residues C-terminal to Scc1’s NTD. Unlike the DL motifs, this motif is highly conserved among eukaryotes (Lee et al., 2016), suggesting that it may have multiple partners. Deletion of the motif *(Δ131-138)* is lethal, as is *scc1V137K,* but we have hitherto assumed that these effects are due merely to loss of Pds5 binding. This may be mistaken. Though *scc1V137K* causes only a modest defect in cohesin loading in cells expressing wild type Scc1 (Fig. S7C), the mutation has a more severe phenotype when cells undergo S phase in Scc1’s absence. Thus, it reduced loading along chromosome arms by twofold and around centromeres by 3.4 fold (Fig. 7E). Equally striking, the cohesin that still loads at *CEN*s fails to translocate normally into peri-centric sequences and instead accumulates in two peaks 300bp on either side of *CEN*s (Fig. 7E). Crucially, neither the loading nor the translocation defect can be attributed to a failure to recruit Pds5 because depletion of the latter under similar conditions, namely 60 min after release from a pheromone-induced G1 arrest, had a much milder effect (Fig. 7F). It had no effect on arm loading and only modest defects in loading around *CEN*s or translocation away from them. Finally, *scc1V137K* caused a modest reduction in tetramer ATPase activity (Fig. S7D), suggesting that it may subtly affect the way Scc2 interacts with cohesin.

In conclusion, we find that Scc2 mediates its effect on cohesin’s ATPase activity as well as loading and translocation in vivo by interacting with possibly three peptide motifs within Scc1 situated between its NTD and its Scc3 binding site. The dramatic effect of V137K on cohesin’s translocation from *CEN*s into peri-centric sequences suggests that translocation is a property intrinsic to cohesin, driven presumably by Scc2’s stimulation of cohesin ATPase activity, and is not mediated merely by an extrinsic motor such as RNA polymerase.

## Discussion

### Scc2 drives cohesin’s DNA-dependent ATPase activity

Unlike Scc3 and Pds5, Scc2 is rarely found stably bound to cohesin and has hitherto been thought of as a separate factor dedicated to loading (Ciosk et al., 2000). The findings described here alter this view radically. We show that Scc2 binds to Scc1 sequences essential for cohesin’s association with and translocation along chromosomes, and that it is necessary for ATPase activity associated with all types of cohesin complexes, be they trimers, tetramers, or hexamers. Remarkably, Scc2 is even sufficient to mediate full DNA-dependent ATPase activity associated with Smc1/Smc3/Scc1 trimers. We conclude that stimulating cohesin’s ATPase in a DNA dependent manner is possibly Scc2’s key function, a process necessary for loading but apparently also for translocation. Our finding that the ATPase activity associated with cohesin trimers is as high as that of tetramers in the presence of DNA and Scc2 implies that Scc3 does not have an essential role in stimulating cohesin’s ATPase, though it may indeed facilitate the process. Scc3’s actual function remains enigmatic.

### Two steps in the loading reaction at *CEN*s

Our observations suggest that the loading reaction at *CEN*s can be divided into two steps and that Scc2 is required for both. The first is formation of cohesin complexes with Scc2 bound to their kleisin subunit in a manner that displaces Pds5 and with ATPase heads engaged in the presence of ATP, while the second is their conversion upon ATP hydrolysis into complexes that stably associate with chromatin and then translocate into neighbouring sequences (Fig. S7E). Notably, this second step is preferentially compromised by mutation within *SCC2* of a pair of highly conserved basic residues (K788 H789) that have a key role in facilitating stimulation of cohesin’s ATPase by DNA. The conferral of DNA-dependent ATPase activity by Scc2 is reminiscent of Ycg1, a condensin HAWK that more closely resembles Scc2 than either Scc3 or Pds5. Indeed, an alignment of Scc2’s structure with that of a Ycg1:DNA co-crystal (Kschonsak et al., 2017) shows that Scc2’s crucial K788 residue corresponds to Ycg1’s R253, which contacts the phosphate backbone, and is also highly conserved.

### Scc2 and Pds5 are mutually exclusive HAWKs

The notion that Scc2 is a bona fide cohesin subunit and therefore a fully-fledged HAWK is novel. It was not previously appreciated because Scc2’s association with cohesin is only fleeting. Association with cohesin at *CEN*s only lasts 1-2 seconds before displacement by Pds5. For this reason, the vast majority of chromosomal cohesin in yeast is associated with Pds5, not Scc2. Nevertheless, we have documented instances of Pds5’s displacement by Scc2, namely at *CEN* loading sites, in cohesin recruited to Scc2/4 tethered at Tet operators, and when Scc2 is added to cohesin tetramers in vitro. Even more striking, the gain of function *scc2E822K L937F* allele displaces a large fraction of Pds5 from cohesin throughout the genome. We suspect that the failure of ChIP-seq to detect chromosomal Scc2 efficiently, either in wild type or even in *scc2E822K L937F* cells, is due to its occupancy of chromosomal cohesin in vivo being briefer than that by Pds5 and therefore difficult to trap with formaldehyde.

Given the reciprocal nature of cohesin’s occupancy by Scc2 and Pds5, one might predict that Pds5 hinders Scc2 binding as well as vice versa. Indeed, Pds5/Wapl reduces the ATPase activity of wild type but not *scc1V137K* tetramers in vitro; Pds5 over-production is lethal to *scc2-45* cells, while Pds5 depletion leads to higher than normal Scc2-dependent loading throughout the genome. We suggest therefore that chromosomal cohesin switches between two states, one with Scc2 bound, which is active as a DNA-dependent ATPase and capable of loading and translocating along chromatin, and another with Pds5 bound, which is largely inactive as an ATPase but is capable of dissociating from chromatin in a Wapl-dependent manner. The finding that *C. thermotolerans* Scc2 orthologs compete for binding Scc1 in vitro (Kikuchi et al., 2016), made independently during the course of our studies, suggests that competition between Scc2 and Pds5 may be a universal feature. Our findings imply that two separate DL motifs are essential for loading mediated by Scc2. Because these are C-terminal to the motif known to bind Pds5, it is currently unclear why Scc2 and Pds5 recruitment is mutually exclusive. As *scc1V137K* adversely affects loading and translocation as well as Pds5 recruitment, we suspect that Scc2 also contacts the Pds5 binding motif. Moreover, it is quite possible that Pds5 binds in addition the more C-terminal sequences necessary for Scc2 binding. Our work suggests that the interchange between Scc2 and Pds5 creates two types of complex, one containing Scc2 capable of loading and another containing Pds5 capable of release.

### Scc2 and Pds5 have unique roles

The notion that cohesin switches between Scc2- and Pds5-bound states helps to explain the enigmatic finding that ATPase head engagement is involved in both loading and release (Elbatsh et al., 2016). Complexes that have already loaded onto chromosomes whose heads have engaged in the presence of Pds5 are capable of releasing DNA (via dissociation of the Smc3-Scc1 interface) in the absence of ATP hydrolysis itself (Beckouet et al., 2016), while heads engaged in the presence of Scc2 at loading sites are capable of entrapping it (possibly via transient dissociation of the Smc1-Smc3 hinge interface) but only after a full cycle of ATP hydrolysis.

Analysis of both loss- and gain-of-function mutations affecting universally conserved surface residues implies that a key function of Scc2/Nipbl is to stimulate in a DNA dependent manner cohesin’s ATPase activity, something that Pds5 apparently cannot perform. Because some of these mutations are associated with CdLS, we suggest that irrespective of whether Nipbl has functions besides regulating cohesin, its ability to control transcription during development depends on its ability to stimulate cohesin’s ATPase. The view of Scc2 and Pds5 emerging from our work is difficult to reconcile with the suggestion that Pds5 and Wapl promote loading in living cells (Murayama and Uhlmann, 2015). As well as being unnecessary for cohesin’s ATPase activity and loading in vivo, Pds5 is actually displaced from complexes engaged in loading. Nevertheless, we cannot exclude the possibility that by temporarily replacing Scc2, Pds5 may facilitate further ATPase cycles mediated by Scc2 and thereby promote cohesin’s translocation along chromosomes.

### Importance of cohesin’s ATPase

Analysis of various *smc1/3, scc1* and *scc2* mutants revealed only a rough congruence between their in vitro ATPase activities and their abilities to load/translocate in vivo. Thus, *smc1E1158Q, smc3E1155Q,* or *scc1 Δ152-256* greatly reduce both cohesin’s ATPase and its loading throughout the genome in vivo, but *scc1V137K* has a profound effect on loading at and translocation from *CEN*s but has only a modest effect, if any effect on ATPase activity, *scc2K788E H788E* abolishes loading but reduces in vitro ATPase activity by only two fold, while the gain of function *scc2E822K L937K* allele increases loading genome wide, especially in the absence of Scc4, and increases ATPase activity by 30%.

Curiously, some of these correlations break down upon addition of DNA; for example, amelioration of the ATPase defect caused by *scc1Δ152-256.* Despite this, it is likely that stimulation of cohesin’s ATPase by DNA is of physiological importance, especially as *scc2K788E H788E* compromises the response. How DNA mediates its effect and how it manages to rescue severely defective mutants is at present unclear. We suggest that at high concentrations in vitro, DNA binds to and stabilizes an “active” conformation whose creation normally depends on other factors. In the case of tetramers, DNA stimulates the Scc2-dependent ATPase merely two fold but in the case of trimers, it increases it by at least ten fold, an effect that is greatly reduced by *scc2K788E H788E.* In conclusion, our data are consistent with the notion that cohesin’s ATPase is crucial for its association with and translocation along chromatin. However, the clear instances where ATPase activity fails to correlate with loading suggest that some if not all of our mutations may compromise biological activity by altering conformational changes in cohesin that are linked to the ATPase cycle as well as the cycle itself.

What these changes are and how they lead to cohesin’s association with and translocation along chromatin are poorly understood. Loading/translocation are normally accompanied by entrapment of DNA inside cohesin rings but this cannot be the sole mechanism by which cohesin loads and translocates because the latter are unaffected by a mutation in cohesin’s hinge domain that greatly reduces entrapment (Srinivasan et al., 2017). Crucially, other mutations in the hinge abolish loading/translocation without affecting cohesin’s ATPase, raising the possibility that ATPase activity driven by Scc2’s association with Scc1 drives changes in the conformation of Smc1/Smc3 hinges that facilitate both loading and translocation, a process that is sometimes but not invariably accompanied by DNA entrapment. Thus, in addition to activating cohesin’s ATPase, Scc2 might help mediate interactions between the hinge and ATPase domains of Smc1/3.

### Scc4 mediates *CEN*-specific loading

Our ability to dissect events that take place during cohesin loading and to distinguish them from translocation was greatly facilitated by the fact that a large fraction of pericentric cohesin is initially loaded at core *CEN*s and only subsequently translocates into neighbouring sequences, up to 30 kb away. We demonstrate here that Scc4 bound to Scc2’s N-terminal sequences has a profound role in this process. Thus, in *scc2E822K L937F* cells capable of proliferating in the absence of *SCC4, CEN*-specific loading is entirely lacking and the pattern of peri-centric cohesin resembles that along chromosome arms, where in contrast to centromeres loading is widely distributed throughout transcription units. Recent observations suggest that Scc4 performs this function by binding directly to the Ctf19 complex (Hinshaw et al., 2017) and thereby greatly increasing at this location high rates cohesin ATPase activity driven by the Scc2. Our bioinformatic analysis of Scc4 and centromere proteins imply that this process co-evolved with that of point centromeres. If pericentric cohesin were also loaded at sites containing CENP-A-like nucleosomes in fungal ancestors with distributed centromeres, then peri-centric cohesin would be loaded at lower rates at numerous locations throughout centromeres. The necessary evolution of specific *(CEN)* loading sites in yeasts related to *S. cerevisiae* created a system whose very high rates of loading at unique sites created a system whose analysis was uniquely tractable.

### Scc2 and loop extrusion

The notion that cohesin’s loading and translocation along chromatin may be driven by cycles of ATP hydrolysis mediated by displacement of Pds5 by Scc2 has important implications for the mechanism of loop extrusion (LE) thought to be responsible for controlling enhancer/promoter interactions during mammalian development. The recent observation that Scc2/Nipbl associates transiently but continuously with chromosomal cohesin in mammalian cells suggests that its function may be to stimulate the ATP hydrolysis needed to drive the translocation along chromatin necessary for LE (Rhodes et al., 2017). Interestingly, Scc2 does not co-localize with cohesin at CTCF sites (Kagey et al., 2010), which have been postulated to block LE (Fudenberg et al., 2016; Sanborn et al., 2015). We therefore suggest that CTCF might block LE by somehow hindering replacement of Pds5 by Scc2, as discussed in a recent bioRxiv pre-print (Wutz et al., 2017). Consistently, depletion of Pds5 in human cells results in a decrease in the definition of TADs and a reduction in the number of loops between convergent CTCF sites (Wutz et al., 2017). Indeed, the exquisite sensitivity of key developmental switches revealed by Nipbl haplo-insufficiency being the cause of Cornelia de Lange syndrome may arise because the precise level of Scc2/Nipbl may determine the rate of ATP hydrolysis and thereby the processivity of LE.

## Author Contributions

N.J.P., T.G.G., and K.A.N. designed and conducted experiments and wrote the manuscript. J.M, B.H., and C.C. designed and conducted experiments. M.V., W.C. and M.S. conducted experiments. B-G.L., and J.L. developed cohesin purification techniques.

## Acknowledgements

We are grateful to Adele Marston for sharing yeast strains, Peter West for programming support, and to Tatiana Wilson, Isobel Johns and Stefanos Skylakakis for invaluable technical assistance. We thank all members of the Nasmyth group for valuable discussions. This work was funded by Cancer Research UK (C573/A12386 to K.N.), the Wellcome Trust (107935/Z/15/Z to K.N.)(202062/Z/16/Z to B.H.), and MRC (MR/L018047/1 to K.N.).

## Materials and Methods

### Yeast strains and growth conditions

All yeast strains were derived from W303 and grown in rich medium (YEP) supplemented with 2% glucose (YPD) at 25°C unless otherwise stated. Cultures were agitated at 200rpm (Multitron Standard, Infors HT). Strain numbers and relevant genotypes of the strains used are listed in the Key Resource Table.

To arrest the cells in G1, α-factor was added to a final concentration of 2mg/L/h, every 30min for 2.5h. Release was achieved by filtration wherein cells were captured on 1.2μm filtration paper (Whatman^®^ GE Healthcare), washed with 1L YPD and resuspended in the appropriate fresh media. To arrest the cells in G2, nocodazole (Sigma) was added to the fresh media to a final concentration of 10μg/mL and cells were incubated until the synchronization was achieved (>95% large-budded cells). To inactivate temperature sensitive alleles, fresh media was pre-warmed prior to filtration (Aquatron, Infors HT).

To arrest cells in late G1 with *GAL-SIC1* arrest, cells were grown in YP supplemented with 2% Raffinose and α-factor was added to a final concentration of 2mg/L/h, every 30min for 2.5h. An hour before release Galactose was added to 2% of the final volume. Release was achieved by filtration wherein cells were captured on 1.2μm filtration paper (Whatman^®^ GE Healthcare), resuspended into YPD, and incubated for 60min at 25°C.

To produce cells deficient of Scc1, the gene was placed under the *MET3*-repressible promoter. Liquid cultures were grown in minimal media supplemented with 2% glucose and 1% -MET dropout solution overnight, diluted to 0D_600_=0.2 and allowed to grow to 0D_600_=0.4. Cells were then collected by filtration as described above, resuspended in YPD supplemented with 8mM methionine and arrested in G1. 0nce arrested, the cells were collected by filtration, washed with YPD in the presence of 8mM methionine and released into the same media.

To produce cells deficient in Pds5 using the AID system, cells were arrested with α-factor as previously described. 30min prior to release, auxin was added to 5mM final concentration. Cells were then filtered as previously described and released into YPD medium containing 5mM auxin.

### Screening for suppressors of *scc4-4*

Forty independent colonies of the parental strain (*smc1D588E*::*TRP1* YCplac33:*scc4-4*::*NATMXscc4Δ*::*HIS3* (K23983)) were picked and grown overnight at 25°C. Each was plated at 5 OD_600_ units per plate over 3 plates and incubated at 35.5°C until colonies appeared. Up to 3 colonies were picked from each plate and streaked for single colonies at 25°C before being retested for growth at 35.5°C. Those that grew at 35.5°C were checked by PCR from genomic DNA preparations for revertants of Scc4. Isolated suppressors that did not show revertant mutations were checked for 2:2 segregation and grouped into complementation groups prior to deep sequencing. To check if for the ability to rescue the deletion of Scc1, suppressors were streaked onto 5-FOA containing medium and allowed to grow for 2 days.

### Protein gel electrophoresis and Western blotting

The samples were mixed with 4X LDS sample buffer (NuPAGE^®^ Life Technologies), loaded onto 3-8% Tris-acetate gels (NuPAGE^®^ Life Technologies) and the proteins separated using an appropriate current. The proteins were then transferred onto 0.2μm nitrocellulose using Trans-blot^®^ Turbo^TM^ transfer packs for the Transblot^®^ Turbo^TM^ system (Bio-Rad).

| <u>Primary antibody</u> | <u>Immunogen</u> | <u>Origin</u> | <u>Dilution</u> |
| --- | --- | --- | --- |
| V5 (Bio-Rad) | PK | Mouse | 1:1,000 |
| 3F10 (Roche) | HA | Rat | 1:5,000 |
| PGK1 | PGK1 | Mouse | 1:5,000 |

For visualization the membrane was incubated with Immobilon^TM^ Western Chemiluminescent HRP substrate (Millipore) before detection using an ODYSSEY^®^ Fc Imaging System (LI-COR).

### Multiple sequence alignment

Multiple sequence alignments were created using Clustal Omega (Sievers et al., 2011).

### Quantitative ChIP-seq

Cells were grown exponentially to 0D_600_–0.5 and the required cell cycle stage where necessary. 15 OD_600_ units of *S. cerevisiae* cells were then mixed with 5 OD_600_ units of *C. glabrata* to a total volume of 45mL and fixed with 4mL of fixative (50mM Tris-HCl, pH 8.0; 100mM NaCl; 0.5mM EGTA; 1mM EDTA; 30% (v/v) formaldehyde) for 30min at RT with rotation. Fixation was quenched with 2mL of 2.5M glycine incubated at RT for 5min with rotation. The cells were then harvested by centrifugation at 3,500rpm for 3min and washed with ice-cold 1X PBS. The cells were then resuspended in 300μL of ChIP lysis buffer (50mM Hepes-KOH, pH 8.0; 140mM NaCl; 1mM EDTA; 1% (v/v) Triton X-100; 0.1% (w/v) sodium deoxycholate; 1mM PMSF; 1 tablet/25mL protease inhibitor cocktail (Roche)) and an equal amount of acid-washed glass beads (425-600μm Sigma) added before cells were lysed using a FastPrep^®^-24 benchtop homogeniser (M.P. Biomedicals) at 4°C (3x 60s at 6.5m/s or until >90% of the cells were lysed as confirmed by microscopy).

The soluble fraction was isolated by centrifugation at 2,000rpm for 3min then sonicated using a bioruptor (Diagenode) for 30min in bursts of 30s on and 30s off at high level in a 4°C water bath to produce sheared chromatin with a size range of 200-1,000bp. After sonication the samples were centrifuged at 13,200rpm at 4°C for 20min and the supernatant was transferred into 700μL of ChIP lysis buffer. 30μL of protein G Dynabeads (Invitrogen) were added and the samples were pre-cleared for 1h at 4°C. 80μL of the supernatant was taken as the WCE and 5μg of antibody (anti-PK (Bio-Rad) or anti-HA (Roche)) was added to the remaining supernatant which was then incubated overnight at 4°C. 50μL of protein G Dynabeads were then added and incubated at 4°C for 2h before washing 2x with ChIP lysis buffer, 3x with high salt ChIP lysis buffer (50mM Hepes-KOH, pH 8.0; 500mM NaCl; 1mM EDTA; 1% (v/v) Triton X-100; 0.1% (w/v) sodium deoxycholate; 1mM PMSF), 2x with ChIP wash buffer (10mM Tris-HCl, pH 8.0; 0.25M LiCl; 0.5% NP-40; 0.5% sodium deoxycholate; 1mM EDTA; 1mM PMSF) and 1x with TE pH 7.5. The immunoprecipitated chromatin was then eluted by incubation in 120μL of TES buffer (50mM Tris-HCl, pH 8.0; 10mM EDTA; 1% SDS) for 15min at 65°C and the collected supernatant termed the IP sample. The WCE extracts were mixed with 40μL of TES3 buffer (50mM Tris-HCl, pH 8.0; 10mM EDTA; 3% SDS) and all samples were de-crosslinked by incubation at 65°C overnight. RNA was degraded by incubation with 2μL RNase A (10mg/mL; Roche) for 1h at 37°C and protein was removed by incubation with 10μL of proteinase K (18mg/mL; Roche) for 2h at 65°C. DNA was purified using ChIP DNA Clean and Concentrator kit (Zymo Research).

### Extraction of yeast DNA for deep sequencing

Cultures were grown to exponential phase (OD_600_=0.5). 12.5 OD_600_ units were then collected and diluted to a final volume of 45mL before fixation as described in the protocol for ChIP-seq. The samples were treated as specified in the ChIP-seq protocol up to the completion of the sonication step whereby 80μL of the samples were carried forward and treated as WCE samples.

### Preparation of sequencing libraries

Sequencing libraries were prepared using NEBNext^®^ Fast DNA Library Prep Set for Ion Torrent^TM^ Kit (New England Biolabs) to the manufacturers instructions. To summarise, 10-100ng of fragmented DNA was converted to blunt ends by end repair before ligation of the Ion Xpress^TM^ Barcode Adaptors. Fragments of 300bp were then selected using E-Gel^®^ SizeSelect^TM^ 2% Agarose gels (Life Technologies) and amplified with 6-8 PCR cycles. The DNA concentration was then determined by qPCR using Ion Torrent DNA standards (Kapa Biosystems) as a reference. 12-16 libraries with different barcodes could then be pooled together to a final concentration of 350pM and loaded onto the Ion PI^TM^ V3 Chip (Life Technologies) using the Ion Chef^TM^ (Life Technologies). Sequencing was then completed on the Ion Torrent Proton (Life Technologies), typically producing 6-10 million reads per library with an average read length of 190bp.

### Data analysis, alignment and production of BigWigs

Unless otherwise specified, data analysis was performed on the Galaxy platform (Giardine et al., 2005). Quality of the reads was assessed using FastQC (Galaxy tool version 1.0.0) and trimmed as required using ‘trim sequences’ (Galaxy tool version 1.0.0). Generally, this involved removing the first 10 bases and any bases after the 200^th^ but trimming more or fewer bases may be required to ensure the removal of kmers and that the per-base sequence content is equal across the reads. Reads shorter than 50 bp were removed using Filter FASTQ (Galaxy tool version 1.0.0, minimum size: 50, maximum size: 0, minimum quality: 0, maximum quality: 0, maximum number of bases allowed outside of quality range: 0, paired end data: false) and the remaining reads aligned to the necessary genome(s) using Bowtie2 (Galaxy tool version 0.2) with the default (--sensitive) parameters (mate paired: single-end, write unaligned reads to separate file: true, reference genome: SacCer3 or CanGla, specify read group: false, parameter settings: full parameter list, type of alignment: end to end, preset option: sensitive, disallow gaps within *n*-positions of read: 4, trim *n*-bases from 5’ of each read: 0, number of reads to be aligned: 0, strand directions: both, log mapping time: false) (Langmead and Salzberg, 2012).

To generate alignments of reads that uniquely align to the *S. cerevisiae* genome, the reads were first aligned to the *C. glabrata* (CBS138, genolevures) genome with the unaligned reads saved as a separate file. These reads that could not be aligned to the *C. glabrata* genome were then aligned to the *S. cerevisiae* (sacCer3, SGD) genome and the resulting BAM file converted to BigWig (Galaxy tool version 0.1.0) for visualisation. Similarly this process was done with the order of genomes reversed to produce alignments of reads that uniquely align to *C. glabrata*.

### Visualisation of ChIP-seq profiles

The resulting BigWigs were visualised using the IGB browser (Nicol et al., 2009). To normalise the data to show quantitative ChIP signal the track was multiplied by the samples occupancy ratio (OR) and normalised to 1 million reads using the graph multiply function.

In order to calculate the average occupancy at each base pair up to 60kb around all 16 centromeres, the BAM file that contains reads uniquely aligning to *S. cerevisiae* was separated into files for each chromosome using ‘Filter SAM or BAM’ (Galaxy tool version 1.1.0). A pileup of each chromosome was then obtained using samtools mpileup (Galaxy tool version 0.0.1) (source for reference list: locally cached, reference genome: SacCer3, genotype likelihood computation: false, advanced options: basic). These files were then amended using our own script ‘chr_position.py’ to assign all unrepresented genome positions a value of 0. Each pileup was then filtered using another in-house script ‘filter.py’ to obtain the number of reads at each base pair within up to 60kb intervals either side of the centromeric CDEIII elements of each chromosome. The number of reads covering each site as one successively moves away from these CDEIII elements could then be averaged across all 16 chromosomes and calibrated by multiplying by the samples OR and normalising to 1 million reads. All scripts written for this analysis method are available on request.

### Identification of mutations from whole genome sequencing

SNPs were called using command line on a local server. First a pileup was created using samtools mpileup (-v --skip-indels -f sacCer3.fa -o *sample name*.vcf *sample name*.bam), then SNPs called using bcftools call (-v -c -o *sample name*.bcf *sample name*.vcf). To find mutations unique to a suppressor strain, lists of SNPs from the parental strain or backcrossed clones of the suppressor strain were compared to the list of SNPs from the suppressor strain. In the case of parental strains, mutations that were present in both were removed and in the case of backcrossed clones of the suppressor strain, mutations that were present in both were kept in order to identify the mutation that caused the suppression phenotype. This was done using ‘MutationFinder.py’ and the resulting lists further narrowed using ‘yeastmine.py’ which searches the Saccharomyces Genome Database (SGD) for genes that correspond to the position of each mutation so that those that lie outside of genes could be removed. From this it was possible to identify the mutation in each suppressor that gave rise to the suppressor phenotype.

### ATPase assay

ATPase activity was measured by using the EnzChek phosphate assay kit (Invitrogen) by following the protocol as provided. Cohesin in various complexes and its subunits was added to a final concentration of 50nM (or as else stated in the main text) and carried out always under 50mM NaCl in the presence of 700nM 40bp dsDNA in those experiments testing the effect of duplex DNA. The reaction was started with addition of ATP to a final concentration of 1.3mM always in a final volume of 150ul. ATPase activity was measured by recording absorption at 360nm every 30s for 1h30min using a PHERAstar FS. ΔAU at 360nm was translated to Pi release using an equation derived by a standard curve of KH_2_PO_4_ provided with the EnzChek kit and according to instructions. The reactions were assumed linear for at least the first 10min of the experiment and rated presented were calculated using this time slot. With the end of the reaction a fraction of each reaction was run in SDS-PAGE and the gel stained with coomassie brilliant blue in order to test that the complexes were intact throughout the experiment and that equal amounts were used when testing various mutants and conditions. At least two independent biological experiments were performed for each experiment.

### In vivo chemical crosslinking

Cells were grown to log phase in YPD, 15 OD_600_ of cells were centrifuged at 3,500rpm at 4°C for 5 minutes. The pellet was washed twice in ice cold PBS, then resuspended in 500μl of EBX buffer (25mM Hepes pH 8, 50mM KCl, 50mM MgSO_4_, 0.25% Triton-X v/v, 0.05% NP-40 v/v, 1/25ml protease inhibitor tablet (Roche), 1mM PMSF, 0.1mg/ml RNase (Roche), Benzonase 1:1000 (Sigma). Each sample was split in two, 12.5μl of 125mM BMOE (in DMSO) was added to one half (+crosslinker) and 12.5μl of DMSO was added to the other (-crosslinker). Samples were incubated on ice for 6 minutes. Acid washed glass beads (425-600μm, Sigma) were added at 1:1 ratio (v/v) and cells lysed using the FastPrep^®^ 824 benchtop homogeniser (M.P. Biomedicals) at 4°C. Lysis was checked by microscopy and the lysates were centrifuged for 10 minutes at 4°C. 700μl of lysis buffer was added with 3μg HA antibody (3f10 Roche) prior to rotation at 4°C for 1hr. 30μl of protein-G Dynabeads (Invitrogen) were added and samples rotated for 1hr at 4°C. The supernatant was discarded and 25μl of 1X Laemmli buffer was added. Samples were boiled at 95°C for 5 minutes, centrifuged for 5minutes, and 15μl of each sample analysed by SDS-PAGE prior to Western blot.

### Recombinant yeast cohesin complex cloning

The Smc1, 8xHis-Smc3, Scc1 2xStrepII, Scc3, Pds5, Wpl1, Scc2 from *S.cerevisiae* were gene synthesized (Genscript, Thermo Fisher Scientific, Epoch Lifescience) to optimize codons for efficient recombinant protein expression. The Smc3 and Scc1 genes were synthesized with N-terminal His and C-terminal StrepII tags, respectively. Flag tag was added into Pds5 gene at the C-termini by PCR. The *SMC1*, 8xHis*SMC3*, *SCC1*2xStrepII, *SCC3*, *PDS5*Flag, *WAPL* genes were inserted into Multibac vectors (pACEBac1, pIDC or pIDS) resulting in the vectors of *SMC1*-pACEbac1, 8xHis*SMC3*-pACEbac1, *SCC1*2xStrepII-pIDC, *SCC3*-pIDC, *PDS5*Flag-pIDS, and *WAPL*-pIDS. Genes in the same Multibac vectors were combined together by Gibson assembly, and *SMC1*-8xHis*SMC3*-pACEbac1, *SCC1*2xStrepII-*SCC3*-pIDC, *PDS5*Flag-*WAPL*-pIDS were generated. The *SMC1*-8xHis*SMC3*-pACEbac1 vector was fused to the *SCC1*2xStrepII-pIDC, *SCC1*2xStrepII-*SCC3*-pIDC, *PDS5*Flag-*WAPL*-pIDS vectors by in vitro Cre recombinase reaction (New England Biolabs), and then the transfer vectors for trimer (*SMC1*-8xHis*SMC3*-pACEbac1/*SCC1*2xStrepII-pIDC), tetramer (*SMC1*-8xHis*SMC3*-pACEbac1/*SCC1*2xStrepII-*SCC3*-pIDC), hexamer (*SMC1*-8xHis*SMC3*-pACEbac1/*SCC1*2xStrepII-*SCC3*-pIDC/*PDS5*Flag-*WAPL*-pIDS) were generated. A similar approach was used for the GFP-ΔN132-*scc2*-1xStrepII bacmid developed. The transfer vectors for trimer, tetramer, and hexamer were transformed into DH10EmbacY cells (Vijayachandran et al, 2011). The isolated bacmid DNAs were transfected into Sf9 cells using Fugene HD reagent (Promega). The generated viruses were infected into Sf9 cells, and the cells were cultured at 27 °C for 72h in Insect-XPRESS protein-free medium with L-glutamate (Lonza).

### Protein purification of the cohesin and Scc2 complexes

All versions of the cohesin complexes purified bear a tween StrepII tag in the *SCC1* kleisin. This is the same for the GFP-ΔN132-Scc2 construct used in this study except the later bears a single Strep-II tag. Typically 500ml of SF-9 insect cells were grown to ~3 million/ml and infected with the appropriate baculovirus stock in a 1/100 dilution. Infection was monitored daily and cells harvested when lethality (assayed by the trypan blue test) reached no more than 70-80%. Cell pellets were then frozen in liquid nitrogen and stored at 80°C. Upon thawing, the pellets were suspended in a final volume of ~65-70ml with HNTG lysis buffer (final concentrations of: 25mM Hepes pH 8.0, NaCl 150mM, TCEP-HCl 1mM and Glycerol 10%) and the suspension was immediately supplemented with 2 dissolved tablets of Roche Complete Protease (EDTA-free), 75μg of RNAse I and 7μl of DNAseI (Roche, of 10U/μl stock). The cells were then sonicated at 80% amplitude for 5s/burst/35ml of suspension using a Sonics Vibra-Cell (3mm microtip). In total 5 bursts were given for every 35ml half of the 70ml suspension (the sonication was always performed in ethanolised ice). A spin at 235,000 x g (45,000rpm on a Ti45 fixed angle rotor) followed for 45 mins. The isolated cleared extract was supplemented with 2mM EDTA and was then used to load a 2x5ml Streptactin column at 1ml/min in an ÄKTA Purifier 100. Wash with HNTG+PMSF 1mM+EDTA 2mM (HNTGPE) followed at 1ml/min to the point of ΔAU_280nm_~0 and protein elution ensued using HNTGPE+20mM desthiobiotin at 1ml/min. Peak fractions were analysed using SDS-PAGE and were further purified in a Superose 6 Increase 10/300 using HNTG as running buffer (free of EDTA/PMSF). The resulting peaks were again analysed using SDS-PAGE and the concentration was determined in Nanodrop using A280. Protein was aliquoted and stocked typically in concentrations ranging from 1 to 3mg/ml.

### Pulldown experiments using holocomplexes

10μg of mouse monoclonal anti-GFP antibody (Roche 1184460001) were coupled to 50μl of of Protein G Dynabeads (Invitrogen) rotating at room temperature for 1hr in a final volume of 200μls per reaction/sample using wash buffer to top up (typically the wash buffer was: Hepes 100mM, NaCl 50mM, Tween 0.04%). The beads were washed twice with 1ml wash buffer using a magnet and finally suspended in 50 μls of wash buffer.

In parallel, ATPase reactions with versions of the holocomplex and (excluding mock reactions) versions of the GFP-ΔN132-Scc2 protein were performed essentially as described elsewhere in Methods (omitting the coupled-enzyme reaction) at 25°C on a benchtop shaker typically for 45 mins. Of the 150μl reaction 1/10th was always kept as input material. The pre-coupled ProteinG-antibody dynabeads were then added (50μls) and the reaction continued for another 45mins (with shaking at 900rpm at room temperature). The beads were then pulled using a magnet and a reciprocal to the input amount was kept as flow-through material. The rest of the flow-through was then discarded and 3x1ml washes ensued using a magnet with the wash buffer at 100mM NaCl (Fig.S7) or 150mM NaCl (Fig.S1). The beads were finally transferred to a new tube and eluted with 50μls of 1X SDS buffer and analysed by SDS-PAGE with coommasie brilliant blue staining.

### Cloning of the GST-Scc2 plasmids and purification of GST-Scc2 and Pds5 of figures 7A and S7A

The cDNA encoding Sc Scc2_171-1504_ was subcloned into pGEX6p-1 that introduced an N-terminal GST tag. The GST-Scc2 plasmid was transformed into *Escherichia coli* strain BL21 (DE3). Protein expression was induced by 0.2mM isopropyl-d-1-thiogalactopyranoside (IPTG) at 20°C overnight. GST-Scc2 was then purified with Glutathione Sepharose 4B resin (GE Healthcare) and stored in the storage buffer (20mM Tris-HCl pH 7.5, 150mM NaCl, 1mM TCEP-HCl) at -80°C. Full-length Sc Pds5 cDNA was subcloned into pFastbacHT vector that introduced an N-terminal His_6_-tag. The Sc Pds5 baculovirus was made with the Bac-to-Bac system (Invitrogen). For protein expression, Hi5 insect cells (Sigma-Aldrich) were infected with the Pds5 baculovirus and cultured for 50hr at 27°C. Cells were harvested, resuspended in buffer I (20mM Tris-HCl pH 7.5, 500mM NaCl and 20mM imidazole), and lysed by sonication. After centrifugation, the supernatant was incubated with Ni^2+^ Sepharose 6 Fast Flow resin (GE Healthcare). The Ni resin was washed with buffer II (20mM Tris-HCl pH 7.5, 1M NaCl, 20mM imidazole), and buffer III (20mM Tris-HCl pH7.5, 100mM NaCl, 20mM imidazole) and eluted with buffer IV (20mM Tris-HCl pH 7.5, 100mM NaCl, 150mM imidazole). The eluted His_6-_Pds5 protein was concentrated and applied onto HiLoad 16/60 Superdex 200 prep grade column (GE Healthcare) that had been equilibrated with buffer V (20mM Tris-HCl pH 7.5, 150mM NaCl, 1mM TCEP-HCl). The purified Pds5 protein was then concentrated to 5mg/ml using an Amicon Ultra-15 centrifugal filter unit (Millipore) and stored at -80°C.

### Protein binding competition assay of *S. cerevisiae* Pds5 and Scc2

The Sc Scc1 _1-566_ and Scc1_1-566_ Δ152-256 cDNAs were subcloned into the pCS2 vector. These plasmids were mixed with a TNT Quick Coupled Transcription Translation System (Promega) and incubated at 30°C for 90min in the presence of ^35^S-methionine. The ^35^S-labeled Scc1 proteins were mixed with Glutathione Sepharose 4B beads bound to 10μg GST-Scc2_171-1504_ in the absence or presence of 10μg Sc*SMC3*head, and incubated for 1 h at 4°C in the binding buffer [20mM Tris-HCl pH 7.5, 150mM NaCl, 0.1% Triton-X100]. After incubation, the beads were washed 4 times with the binding buffer. The bound proteins were separated on SDS-PAGE gels, which were stained with Coomassie blue, dried and analyzed with a phosphorimager (GE Healthcare).

For the Pds5 competition assay, Sc Scc1_1-256_ was subcloned into the pCS2 vector and translated *in vitro* with the TNT Quick Coupled Translation System (Promega). The ^35^S-labeled Scc1_1-256_ protein was incubated with varying concentrations (1.2μM, 6.0μM) of Sc Pds5 for 2h at 4°C in 50μl of the binding buffer [20mM Tris-HCl pH 7.5, 150mM NaCl, 0.1 % Triton-X100] in the presence of *SMC3*head. After incubation, the protein mixture and GST-Scc2 were added together to Glutathione Sepharose 4B beads. The reaction mixtures were further incubated for 1 h at 5°C. The beads were washed 4 times with the binding buffer. The bound proteins were separated on SDS-PAGE gels, which were stained with Coomassie blue, dried and analyzed with a phosphorimager (GE Healthcare).

